# Bayesian analysis and efficient algorithms for single-molecule fluorescence data and step counting

**DOI:** 10.1101/2025.03.04.641510

**Authors:** Chiara Mattamira, Alyssa Ward, Sriram Tiruvadi Krishnan, Rajan Lamichhane, Francisco N. Barrera, Ioannis Sgouralis

## Abstract

With the growing adoption of single-molecule fluorescence experiments, there is an increasing demand for efficient statistical methodologies and accurate analysis of the acquired measurements. Existing analysis frameworks, such as those that use kinetic models, often rely on strong assumptions on the dynamics of the molecules and fluorophores under study that render them inappropriate for general purpose step counting applications, especially when the systems of study exhibit uncharacterized dynamics. Here, we propose a novel Bayesian nonparametric framework to analyze singlemolecule fluorescence data that is kinetic model independent. For the evaluation of our methods, we develop four MCMC samplers, ranging from elemental to highly sophisticated, and demonstrate that the added complexity is essential for accurate data analysis. We apply our methods to experimental data obtained from TIRF photobleaching assays of the EphA2 receptor tagged with GFP. In addition, we validate our approach with synthetic data mimicking realistic conditions and demonstrate its ability to recover ground truth under high- and low-signal-to-noise data, establishing it as a versatile tool for fluorescence data analysis.

**Significance statement:** Protein complexes are critical for cell function. Advances in fluorescence experiments have facilitated their direct study, achieving single-molecule resolution in the determination of their stoichiometry. However, the analysis of raw fluorescence data remains a challenge. Traditional visual inspection methods are often time-consuming and susceptible to user bias, while conventional statistical approaches are limited by strong assumptions regarding molecular and probe dynamics or exceedingly high computational requirements. In this study, we introduce a novel statistical methodology for robust and efficient analysis of fluorescence data. Our innovative approach not only enhances the accuracy and reliability of data analysis following a fluorescence experiment but also allows high-throughput applications. Our methods present a significant advancement that may extend the scope of current fluorescence techniques.

## 1 Introduction

The fundamental functions of living cells are carried out or regulated by proteins that assemble in complexes of different sizes [1–3]. Determining the composition of these complexes is crucial for understanding their biological function [4–6]. In recent decades, several fluorescence-based methodologies have been developed to determine protein stoichiometry [7–11], with photobleaching step analysis (PBSA) being one of the most widely used [12, 13]. PBSA relies on the labeling of each molecule of interest with a fluorophore that, during an experiment, initially fluoresces and subsequently photobleaches and stops emitting light within seconds to minutes of the onset of the experiment [14]. Photobleaching events are recorded and used to determine the number of fluorophores within a diffraction-limited fluorescent spot by counting the number of drops in fluorescence intensity over time [15, 16].

The classification of the fluorescence traces obtained from the photobleaching assays is often performed by visual inspection [17–19]. This is time-consuming and, most importantly, subject to human bias. Moreover, single-molecule fluorescence traces are contaminated with noise from multiple sources: photon shot noise [20, 21], camera readout noise [22], and optical drift [23]. This results in a large portion of the traces being excluded from visual classification due to their low signal-to-noise ratio (SNR), which leads to additional bias and uncertainty.

Characteristically, the analysis of the data we use in this work, which was acquired using a novel single-molecule pulldown-POP technique (SiMPull-POP) for the TIRF photobleaching assay [19], is currently limited by reliance on user choices, low SNR, and time constraints, among other challenges [24–29]. In this assay, proteins of interest with a fluorescent probe or protein are immobilized within a sample chamber and imaged by total internal reflection fluorescent (TIRF) microscopy. This method provides single-molecule resolution of individual fluorescent intensities over time that can be used to visualize distinct photobleaching events. In turn, this information provides insight into the stoichiometry of protein assemblies, furthering our understanding of how proteins interact and alter cellular behavior.

Several statistical methodologies have been developed to address the challenges and limitations of time series analysis of single-molecule data [16, 30–43]. The work on molecular quantification in photoactivation localization microscopy (PALM) [34] using a stochastic approach adapted from aggregated Markov methods provides a prototypical methodology for molecular counting problems. However, its scalability and applicability to very large datasets are limited by the reliance on noise assumptions and computational complexity. Recently, a more comprehensive Bayesian nonparametric method was developed implementing specialized Monte Carlo algorithms [35]. This study serves as a reference for determining both the number of fluorophores and their individual photophysical state trajectories. However, it does present a computational bottleneck when dealing with high molecular counts. Moreover, similarly to other Hidden Markov Model (HMM) frameworks for single-molecule time series analysis [36–38, 44], it relies on the assumption that the system in question evolves in a discrete state space and the state-to-state transitions follow *Markovian dynamics*. Markovianity of measurements is equivalent to exponential duel times [45, 46]. Unfortunately, this is a critical assumption that can be violated, and for this reason various HMM methodologies that employ state aggregates have been proposed [47, 48]. In addition, most often, the transition rates are assumed to be slow compared to the data acquisition rate, set by the frame rate and exposure period of the cameras used. A Hidden Markov Jump Process (HMJP) approach for fast kinetics was developed to address this limitation [39, 40]. However, despite using weaker assumptions on the data acquisition timescale, HMJP maintains strong kinetic assumptions of the molecules under study. An alternative approach that relaxes the kinetic assumptions is offered by model-independent methods that do not rely on motion models. We can only find one such example in the literature [41] where a generic step detection algorithm is applied and the computations for step identification are carried out. However, this work is limited by the sensitivity to noise levels and step sizes, and computations rely on likelihood ratios with a fixed threshold selection requiring manual user input.

In this study, we develop a new, computationally efficient, model-free Bayesian nonparametric methodology for the analysis of single-molecule fluorescence data and PBSA. Specifically, we provide a novel statistical framework for the analysis of raw PBSA data and propose, for its evaluation, four computational algorithms with increasing complexity. Our first two algorithms are formulated by combining existing Markov Chain Monte Carlo (MCMC) sampling techniques commonly used within the nonparametric Bayesian framework. The third utilizes a novel sampling strategy that aims to address the shortcomings of the first two algorithms and increase the efficiency and reliability of the parameter estimates. Lastly, additional improvements are made through a fourth algorithm, whose performance is largely independent of the MCMC fine-tuning. We also apply our algorithms to experimental datasets obtained using a novel SiMPull-POP technique for TIRF photobleaching assays and validate them using synthetic data with high and low SNR levels. Finally, we compare the quality and efficiency of the four samplers by assessing their ability to recover the ground truth from synthetic data that mimic realistic laboratory conditions.

The remainder of our study is structured as follows. In Results, we demonstrate and validate our algorithms. In Discussion, we describe the general benefits of our sampler and its wider potential for step counting by photobleaching data analysis. Lastly, in Methods, we present our data analysis and acquisition methods, with additional details provided in Supporting Materials.

## 2 Results

We first apply our methods to analyze single-molecule fluorescence data from EphA2-GFP experiments obtained in TIRF photobleaching assays paired with the recently developed SiMPull-POP technique [19, 24]. Next, we validate our methods using data from simulated experiments and demonstrate their ability to accurately recover the ground truth from traces of both high and low signal-to-noise ratios. Then, we compare the performance of our most advanced MCMC sampler with that of the three simpler ones described in Methods and demonstrate that its complexity is essential for its high performance. Lastly, we compare our most advanced algorithm with existing ones commonly used for fluorescence data analysis and assess its quality under a wide range of experimental conditions demonstrating that it is a reliable alternative for general applications with broader scope and lower computational demands.

To facilitate visual comparison of our results with the data under analysis, in the following, we display measurements *w*_*n*_ and signals *U* (*t*) after rescaling them as described in appendix C. Our rescaling is essentially a linear transformation that standardizes their units, allowing for a direct scale-free comparison between them without altering their characteristics.

### 2.1 Demonstration with laboratory data

To demonstrate the ability of our methods to analyze experimental data and recover the total number of steps *B*, the background intensity *c*_bck_, the intensities of the steps 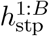, and the times at which the steps occur 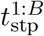, we apply them on raw fluorescence traces [19]. Initially, we choose example data sets with a strong step signature so that they can be easily reproduced with synthetic data when validating the samplers later on. Examples with weaker step signatures are given in the next section and in section 3.2, where results can be compared directly to ground truth values.

Following common practices, we configure our sampler with the default hyperparameters and run it for 10, 000 iterations. From the generated samples, we discard the first 20% due to MCMC burnin [49–51]. We then apply the relabeler described in appendix A.1 to ensure the temporal ordering of the steps. The specific values of the hyperparameters used can be found in appendix E. Similarly to all Bayesian approaches, the hyperparameters can potentially affect the outcomes. In order to demonstrate that this is not the situation in our case, we present a comprehensive sensitivity analysis in appendix F.

The results are shown in fig. 2 and fig. 3. Figure 2 shows three characteristic traces with one, two, or three steps along with their corresponding maximum a posteriori (MAP) estimator of the signal, which is the signal with the highest probability according to our model. The number of steps identified by the MAP signal correlates with the stoichiometry of the protein complexes under study [19]. The arrows highlight photobleaching events. As a sanity check, we also analyzed a trace extracted from a region with no fluorophores; see fig. S2. To ensure the absence of fluorophores, we selected a ROI from the right channel, which exclusively detects red fluorophores and thus remains dark under our experimental conditions. As expected, our sampler correctly identified no steps.

**Figure 1.**
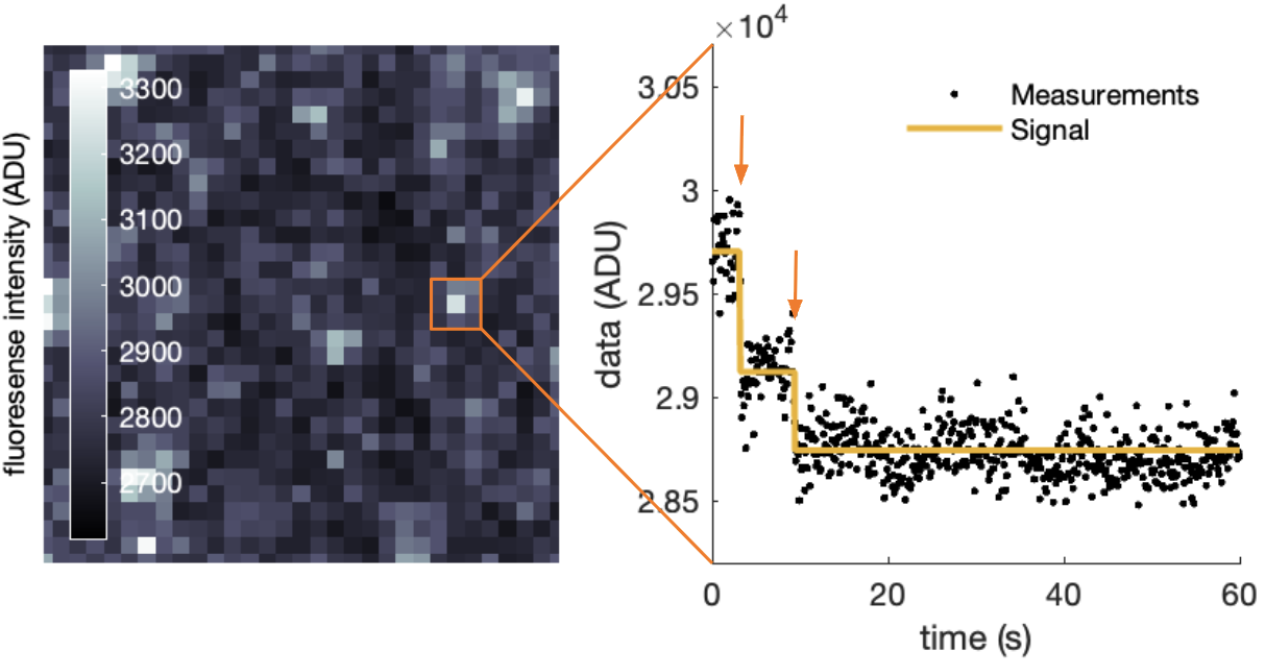
Photobleaching data and step analysis. The panel on the left shows an example of raw fluorescence data obtained from a TIRF microscopy experiment. Lighter pixels correspond to higher fluorescence levels, which indicate the presence of active fluorophores. For each pixel, we can extract the corresponding fluorescence intensity over time for a set region of interest (ROI). In this example, we sum the fluorescence intensities of all nine pixels, each of size 0.16 µm, within the chosen 3×3 ROI collecting photons from a total area of 0.23 µm^2^. The extracted time series is shown in black on the right panel. Then our novel Bayesian framework can be applied to determine the MAP signal estimator (yellow line), provide the total number of steps present, and characterize their time (arrows).

**Figure 2.**
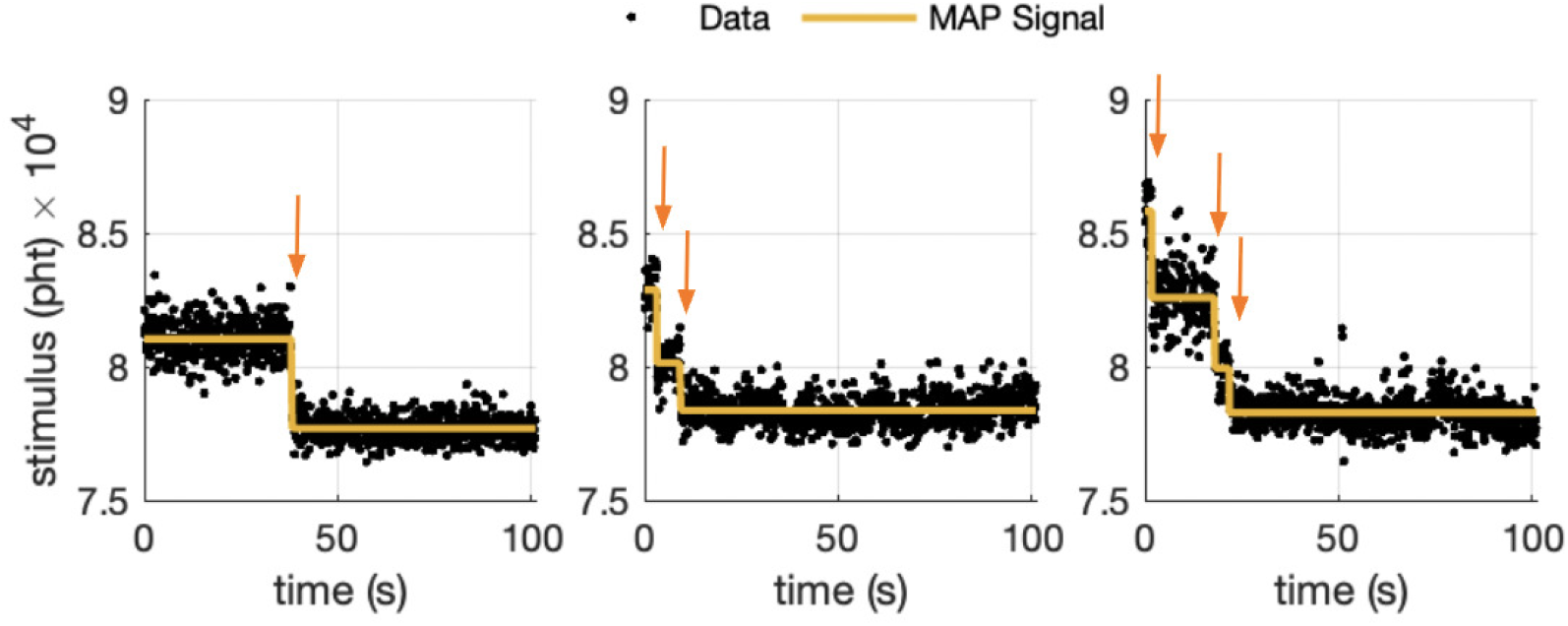
EphA2-GFP data analysis results for a one-, two-, and three-step traces. Experimental fluorescence reduced data are shown in black, and the MAP signal obtained from applying our most advanced sampler is shown in yellow. Arrows identify the estimated photobleaching events.

**Figure 3.**
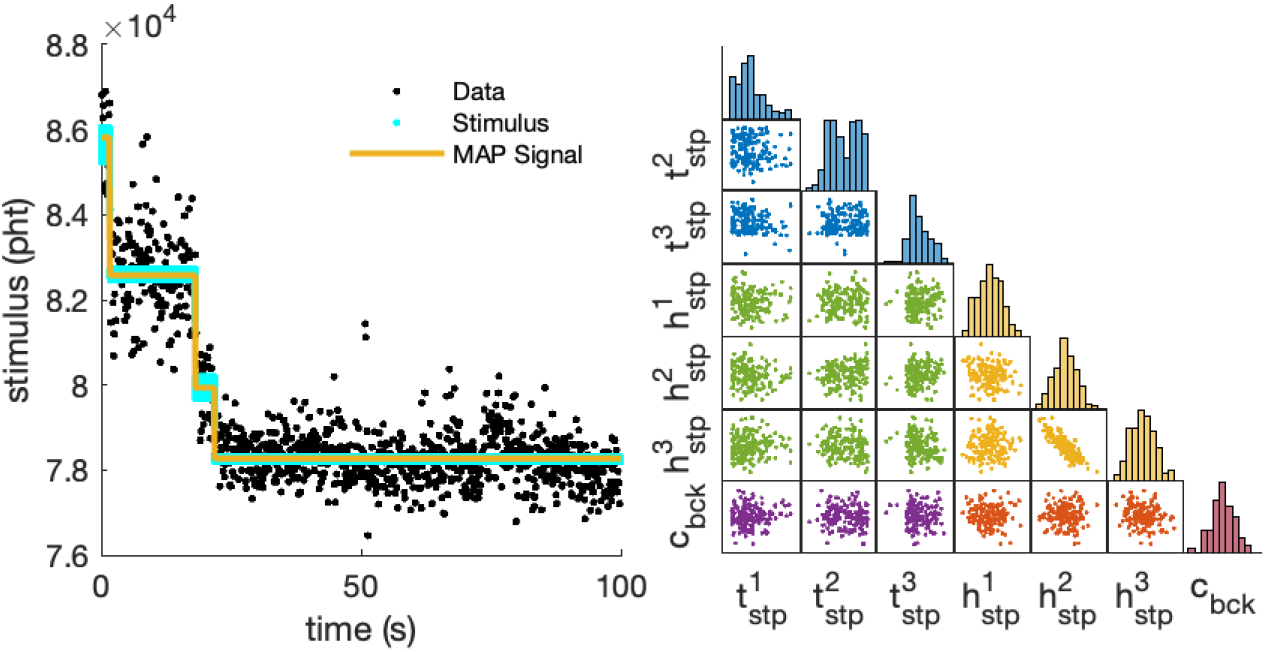
Laboratory data analysis results. On the left panel, we show the experimental fluorescence reduced data (black) along with the stimuli (cyan) at each measurement time t_n_ for each sampler iteration after the burn-in period and the MAP signal (yellow) obtained from applying our sampler. On the right panel, we plot histograms of the sampled 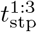 (blue), 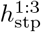 (yellow), and c_bck_ (red) after running 10, 000 iterations and discarding the first 2, 000 as well as scatter plots to show the correlation between variables. Thinning was applied to better showcase the density of the plots.

In fig. 3, we show in more detail the experimental data set for the three-step trace, together with the estimated stimulus traces and the signal MAP estimator. We also show a summary of the posterior probability distribution of each variable of interest and the correlation among them. This posterior propagates uncertainty from the measurements, induced by noise and gaps between time points, and provides not only the best choices, but also confidence intervals around our estimators [46, 52].

As we see in fig. 3, our methods detect 3 steps (left panel). In addition, our methods characterize these steps by estimating the intensity 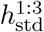 and time 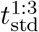 of each step, and background *c*_bck_ (right panel). In particular, the characterization includes estimates of the values and their uncertainly (histograms) as well as correlation between them (scatter plots). While no noticeable correlation between most variables is detected; as expected, there is a strong negative correlation between some of the intensities of consecutive steps. In this example, negative correlation is most prominent between 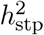 and 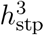 and indicates that small estimation errors in the value of one intensity cause an opposing error in the value of the preceding step’s intensity. In particular, the estimation of 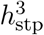 is weaker due to the short lifetime of the 3^rd^ step. Because of the additive nature of our signal, this uncertainty is transmitted to 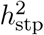 which needs to be negatively adjusted so the total signal level right before the time of the 2^nd^ step matches with the observed data. Similar negative correlations are less prominent between 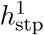 and 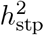 due to the long lifetime of the 2^nd^ step and also between 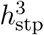 and *c*_bck_ due to the long lifetime of the background.

### 2.2 Demonstration with simulated data

#### 2.2.1 Sampler validation

To validate the reliability of our analysis, we apply it to synthetically generated time traces that mirror the experimental data used in section 2.1. Hence, we prescribe the total number of steps to 3 which recreates the photobleaching events that take place in the experimental data displayed in fig. 3. In addition, we set the timing and camera parameters according to the settings used to collect the measurements as described in Methods. Lastly, we prescribe step intensities, times, and background to the MAP estimators obtained from the analysis of the experimental data; that is, we set the ground truth values of our variables to closely match the experimental data set. In this way, we achieve high fidelity between our simulated experiments and those in the laboratory.

The resulting estimates are shown in fig. 4 together with the data, ground truth, and MAP signal (left panel). The close agreement between the ground truth and MAP signal showcases our methods’ ability to recover the true values for all quantities of interest. A closer look at how the true value of each individual variable compares with the generated posteriors can also be seen in fig. 4 (right panel). For all estimated variables, the ground truth is close to the highest posterior probability values and, as shown in table 1, the bias is minimal across all variables. This quantitatively indicates that our sampler identifies the ground truth with high accuracy.

**Table 1:** Comparison of ground truth values and mean values of generated samples of 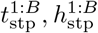, and c_bck_ for both low and high SNR data. Bias is calculated by taking the difference between mean values and ground truth values.

| Scenario | High SNR |  |  | Low SNR |  |  | Units |
| --- | --- | --- | --- | --- | --- | --- | --- |
| Parameter | Ground Truth | Sample Mean | Bias | Ground Truth | Sample Mean | Bias |  |
| $t_{\text{stp}}^1$ | 1.69 | 1.68 | −0.01 | 1.69 | 1.68 | −0.01 | s |
| $t_{\text{stp}}^2$ | 18.12 | 18.13 | 0.01 | 18.12 | 18.20 | 0.08 | s |
| $t_{\text{stp}}^3$ | 21.90 | 21.89 | −0.01 | 21.90 | 22.11 | 0.21 | s |
| $h_{\text{stp}}^1$ | 30.30 | 31.55 | 1.25 | 6.06 | 6.56 | 0.05 | pht/ms |
| $h_{\text{stp}}^2$ | 26.70 | 26.50 | −0.20 | 5.34 | 4.97 | −0.37 | pht/ms |
| $h_{\text{stp}}^3$ | 16.80 | 16.70 | −0.10 | 3.36 | 3.96 | 0.60 | pht/ms |
| $c_{\text{bck}}$ | 783.00 | 782.93 | −0.07 | 783.00 | 782.96 | −0.04 | pht/ms |

**Figure 4.**
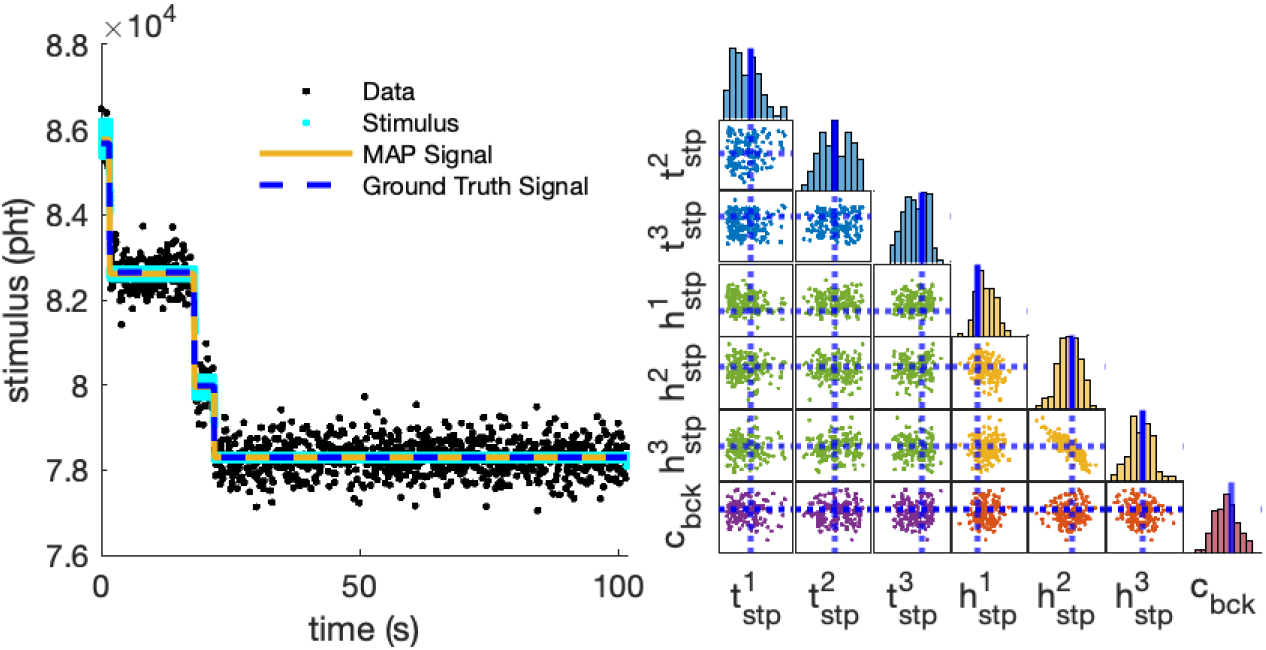
High SNR synthetic data analysis results. On the left panel, we show the reduced synthetically generated fluorescence data (black) and summary results obtained from applying our sampler. Specifically, the stimulus at each measurement time t_n_ for each sampler iteration after the burn-in period is shown in cyan, the MAP signal is shown in yellow, and the ground truth signal is shown as a red dash line. On the right panel, we can see a comparison between sampled values for 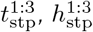 and c_bck_ and true values (blue lines). A total number of 10, 000 iterations was ran and the first 20% were discarded. Thinning was applied to better showcase the density of the scatter plots. A quantitative comparison between sampled and predicted values is shown in table 1.

To further assess our sampler performance on identifying the correct number of steps along with the correct step characteristics, we consider additional synthetic data sets with low signal-to-noise ratio (SNR) mimicking more demanding experimental conditions. We generate these by reducing all step intensities to 1*/*5 of those in the previous example, and maintain all other parameters unchanged. Unlike the higher SNR scenario, which to a certain degree has visually distinguishable steps, the resulting data have a weak signature that is difficult to distinguish from the background. Thus, it cannot be robustly analyzed through visual inspection and instead has to rely on statistical analysis.

Applying our method to this dataset produces the results shown in fig. 5. In this figure, we highlight the estimated times of potential photobleaching events with vertical lines. The density of these lines reflects the certainty of the corresponding photobleaching time. For instance, the first photobleaching event (marked by vertical blue lines) has a strong signature, indicating high certainty in its timing. In contrast, the final step (green lines) has a weaker signature, suggesting greater uncertainty in its timing.

**Figure 5.**
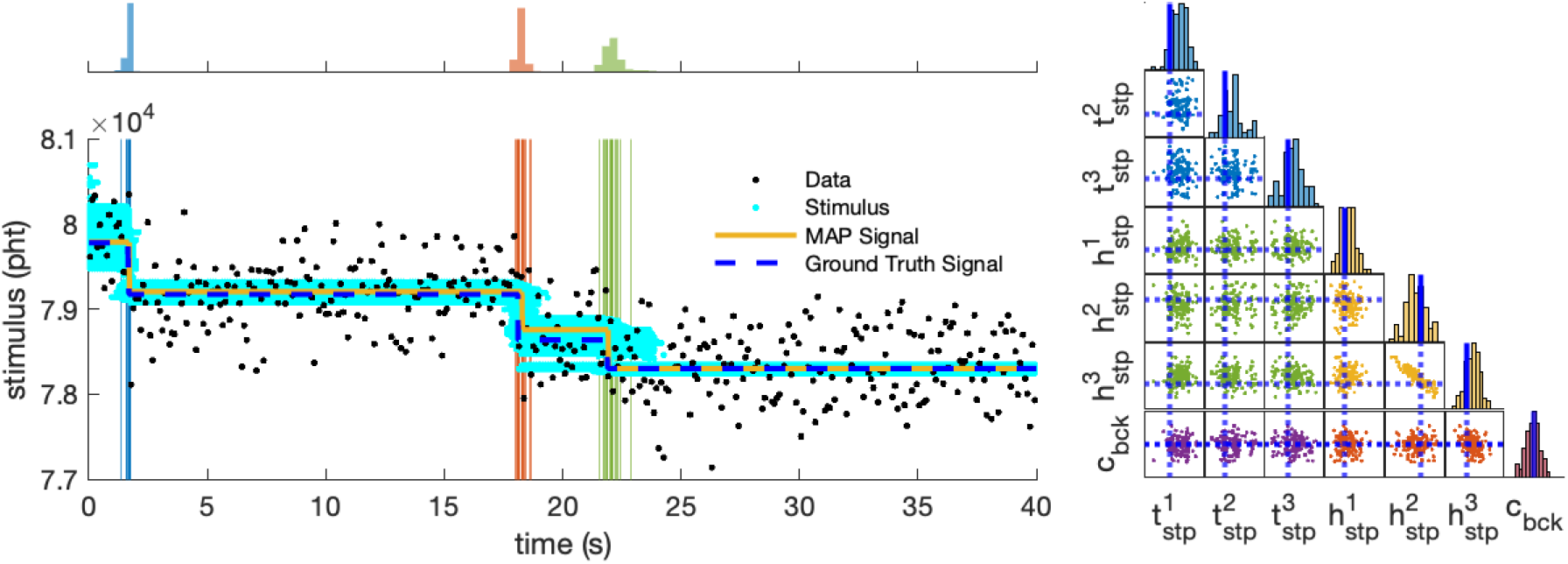
Low SNR synthetic data analysis results. The bottom left panel illustrates the generated data, ground truth signal, MAP signal, and stimuli obtained from our analysis. In the top left panel, we plot histograms of the sampled times of the active steps through the signal’s time course. For clarity, we only plot the first 40 s, which contain all steps. The right panel shows a comparison between sampled values and ground truth of 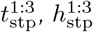 and c_bck_ as well as histograms and scatter plots between them.

To aid in interpretation, we also show histograms of possible photobleaching event times at the top. Here, the spread of a histogram represents uncertainty on the timing, while its integration (i.e., the area under the histogram) indicates the likelihood that the step occurs. Although the estimated stimulus trace shows a higher degree of variation compared to the high SNR scenario, the overall MAP signal still closely follows the true signal. This result demonstrates our methods’ ability to accurately recover the ground truth for our target variables, even in cases where the SNR is exceedingly low. A quantitative comparison between the true and predicted values is shown in table 1, where we see that the difference between the mean of the sample and the ground truth remains small.

#### 2.2.2 Samplers comparison

Thus far, we demonstrated our method’s ability to characterize raw experimental data. For these we applied the fourth sampler described in Methods which is the most advanced mathematically. In the following, we compare the performance of all samplers by applying each to a synthetically generated data set and evaluating the degree to which the ground truth is recovered. Moreover, we assess their ergodicity and mixing times. Both are crucial characteristics for reliable data analysis in practical applications.

For consistency, we apply each sampler to the same synthetic data generated in section 2.2.1 that mimic the data used for the experimental demonstration in section 2.1. The results are shown in fig. 6. As showcased by the mismatch between the ground truth and MAP signal, sampler 1 fails to correctly identify the underlying ground truth step locations and intensities. Moreover, both samplers 1 and 2 overestimate the total number of steps, which in turn would lead to incorrect conclusions regarding the stoichiometry of the molecule under study. Notably, both samplers incorrectly place a step towards the end of the collecting time window.

**Figure 6.**
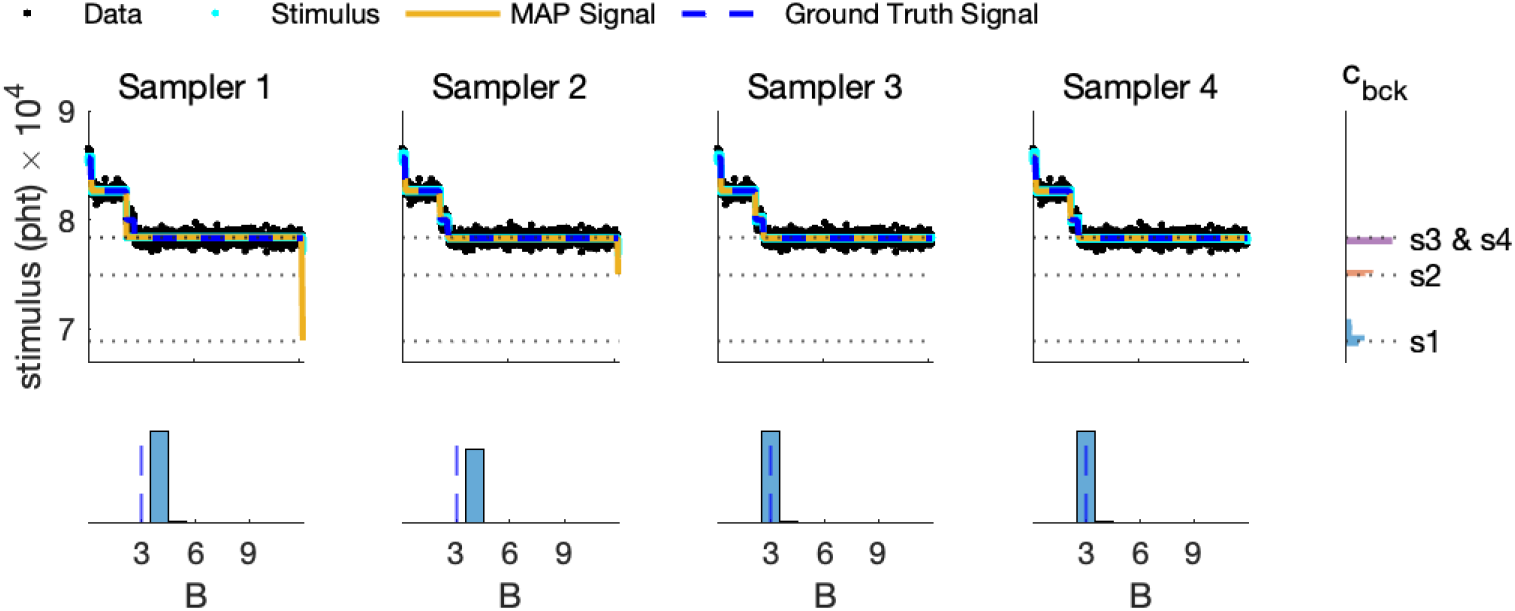
Samplers comparison with synthetic data. The first four panels display the generated data, stimuli, MAP and ground truth signal for each sampler. Underneath each one, we plot histograms for their respective total number of active steps. The top right panel shows histograms for the background samples c_bck_ for each sampler (s1-s4).

The failure of samplers 1 and 2 to properly characterize our model’s posterior is attributed to the nonidentifiability of some of the model parameters, which, in turn, gives rise to multiple local maxima in our probability landscape. For example, different combinations of the background *c*_bck_ and the step intensities 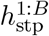 can lead to similar stimuli, which paired with non-informative priors result in multiple local maxima in the parameter space. To overcome this challenge, a sampler needs to have good mixing so that it can efficiently explore the entire parameter space without being trapped in the local maxima [49]. Samplers 1 and 2, by updating only one variable at a time, once they reach a local maximum, cannot cross regions of low posterior probability to reach a different high-probability region [49]. Conversely, by making a simultaneous update of multiple parameters, samplers 3 and 4 are able to efficiently explore the parameter space along multiple axes. In particular, performing a stochastic update on one variable and a simultaneous deterministic update on another allows us to direct these samplers towards areas of high probability, as described in more detail in Supplement. Samplers 3 and 4 are not only able to identify the correct number of steps, location of the steps, and intensity, but also able to quickly reach the correct solution without getting stuck.

To further assess the mixing properties of each sampler and therefore the overall computational time needed, we obtain the autocorrelation function for the background variable *c*_bck_ for each sampler, which we plot in fig. 7. We choose *c*_bck_ because this is the only parameter in our model not affected by label switching [46], so our empirical assessment of the autocorrelation function is more robust. The slow initial decay towards zero along with the overall pattern of the autocorrelation functions of samplers 1 and 2 indicate a significant degree of correlation between samples. However, the autocorrelation functions of the more advanced samplers present a fast initial decay and settle around zero thereafter, indicating that there is little to no relationship between the samples. This means that the overall computational time required to generate a sufficiently large number of samples is significantly lower for samplers 3 and 4, speeding up the analysis and allowing more traces to be analyzed. We also calculate the autocorrelation time, which we define as the first lag where the absolute value of the autocorrelation function falls below 0.1 and therefore indicates how quickly the samples become decorrelated. The autocorrelation times of samplers 1, 2, 3, and 4 are 2120, 845, 52, and 2 lags, respectively. This highlights sampler’s 4 high efficiency and rapid mixing that renders it the sampler of choice for general applications, especially those involving high-throughput data analysis.

**Figure 7.**
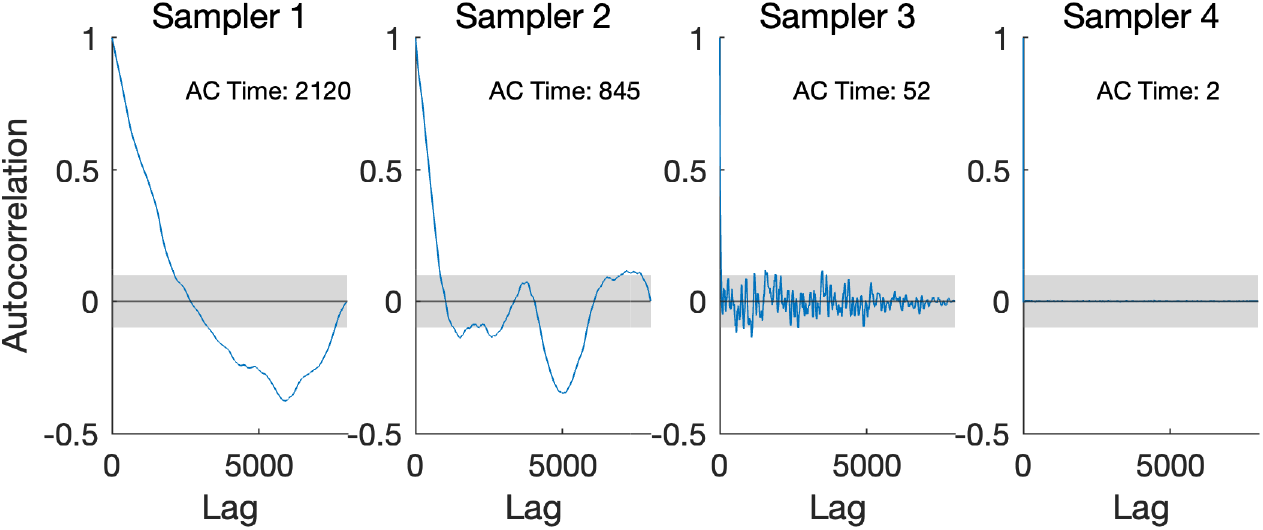
Plots of the autocorrelation function (ACF) of the background parameter c_bck_ sampled from each of the four samplers. Autocorrelation times are displayed in the top right corner of each plot. The number of lags used corresponds to the number of samples kept after the burn-in period.

## 3 Scope assessment

Having demonstrated the validity of our approaches to the successful analysis of SimMPul-POP data, we now provide characteristic results indicating its applicability to more general photobleaching scenarios.

### 3.1 Performance comparison with existing methods

Existing analysis methods that allow for step finding can be classified into two families: model-specific and model-agnostic. Methods in the *first family* make strong assumptions about the dynamics as well as signal formation and noise statistics of the fluorescent system at hand. The most prominent members in this family are methods developed by means of hidden Markov models (HMM) and its variants [36–40, 44]. In contrast, the *second family* methods make no assumptions about dynamics, signal, and noise. Instead, they apply information criteria, such as BIC, and seek to identify and subsequently characterize steps only based on abrupt variations in signal levels and noise statistics without imposing a particular relationship between step times or the statistics of the measurements at different signal levels. The methods in this family are generic analysis tools that can also be applied on non-fluorescence measurements and most prominent members include the methods in [41, 53, 54].

Our methods lie between the two families. On the one hand, we have dedicated noise and signal representations that are specific to fluorescence data. For example, our model accounts for shot-noise caused by stochastic photon detections, excess noise caused by the camera read-out electronics, and realistic data quality curves in agreement with EMCCD and CMOS cameras, for details see appendix A.1. Such specialized representations help to obtain improved inference with low-SNR data relative to the methods in the second family, because our model structure allows only for estimation of parameters that are physical and remain meaningful within a narrow context. On the other hand, we adopt no representation of dynamics or kinetic schemes. This allows our estimated step times to occur equally well near the beginning, middle, or end of a fluorescence trace without having to adhere to dwell times of a set distribution, for details see appendix A.3. This helps inference with data of uncharacterized photodynamics relative to the methods in the first family, because it can avoid statistical bias due to imposed exponential dwell times which are implicit to Markov models [46]. Additionally, our method allows for steps that occur in a *continuum* of times over the signal’s time course, including times between successive measurements or within exposure windows, in stark contrast to the discrete times allowed by HMM. This characteristic helps our method avoid artifactual steps caused by photobleaching events happening mid-through an exposure window. Finally, our combination of agnostic dynamics with dedicated signal and noise representations leads to improved computational performance and fast run-times relative to the methods in both families, as they allow for the development and application of specific computational methods such as those in our most advanced MCMC sampler, for details see appendix B.4. In particular, our computations avoid forward-filtering or greedy evaluations, such as in HMM [44, 47, 48] and Kalafut [41] that create computational bottlenecks in these methods.

To demonstrate important key advantages of our method to HMM based approaches we compare them head-to-head on two example traces. We produce these traces synthetically so that we maintain the ground truth for unambiguous testing. In addition, to facilitate clarity of the presentation, we opt for traces of relatively low noise.

In the first trace, see fig. 8, we simulate a single step occurring mid-though an exposure window. Due to the integrative nature of light detection [39], where photon contributions are accumulated during an entire exposure and converted to a single numerical measurement, this trace contains data at three different levels: one corresponding to the step’s signal, one corresponding to the background, and an artifactual middle level that receives only a single data point near the average of the signal and background levels. Because our method relies on realistic signal formation assumptions, for details see appendix A.1, our method isolates the artifactual middle measurement and correctly identifies the right number of steps. In contrast, due to the fundamental assumption of discreteness of time [46], HMM overestimates the number of steps by introducing an artifactual step to accommodate the middle data point. Such an overestimation is an important limitation of HMM in single-molecule assays, as exposure times are typically long, and therefore the chances of encountering artifactual measurements at the photobleaching times are high as well.

**Figure 8.**
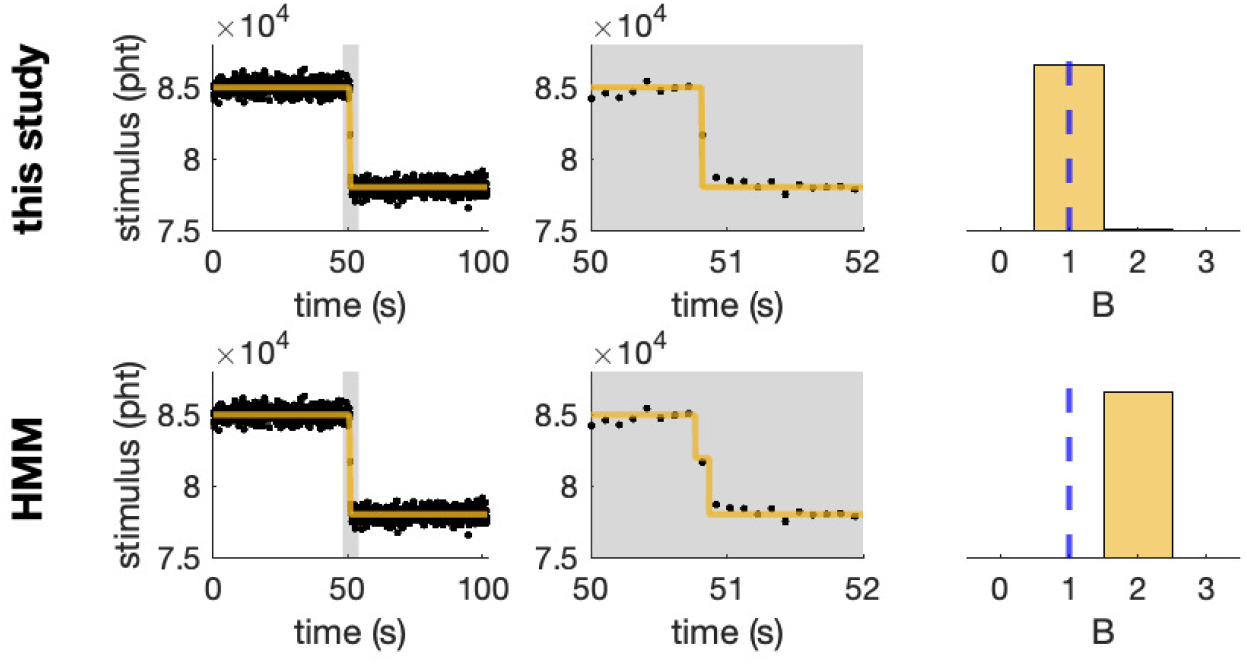
First comparison between HMM and our method. In the leftmost panels, we show a synthetic dataset (black) alongside the MAP signal generate by the specified method (yellow). The middle panels present a zoomed-in view around the photobleaching step. In the rightmost panels, we plot histograms of the number of steps, with the ground truth (vertical line) overlaid for reference.

In the second trace, see fig. 9, we simulate a ladder of steps. This time we compare our method with a Bayesian formulation of the HMM that relies on the common priors mediated by Dirichlet distributions [42, 44, 55]. Furthermore, we intentionally configure our HMM’s hyperparameters so that we induce a strongly informative prior that influences photobleaching dynamics of short dwell times. Although HMM can now correctly estimate the number of steps, as can be seen, it does so with considerably less confidence than our method. Furthermore, due to the strong prior on the dynamics, the reconstructed signal is biased towards noticeably shorter photobleaching times that have little resemblance with the actual measurements. Such dependence of the Bayesian HMM on its specified hyperparameters is a limitation particularity important for applications in step counting, which require implementation of HMMs of a varying number of parameters. In turn, such HMM are achieved only through priors similar to our setup. Unluckily, configuring the priors with the appropriate choice of their hyperparameters requires knowledge of the underlying dwell time-scales that, most often, is unavailable.

**Figure 9.**
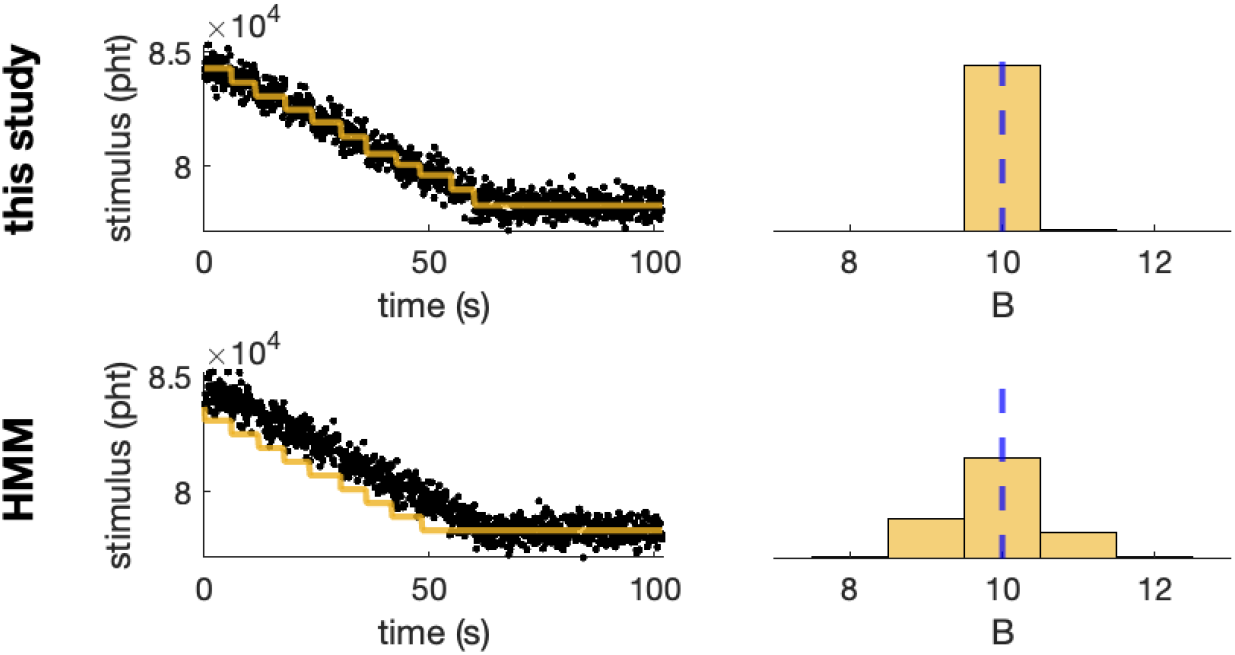
Second comparison between HMM and our method. In the left panel, we show a synthetic dataset (black) alongside the MAP signal generate by the specified method (yellow). In the right panel, we plot histograms of the number of steps, with the ground truth (vertical line) overlaid for reference.

To demonstrate the computational advantages of our method to greedy based approaches we compare them head-to-head on synthetic traces of varying size. In our comparison, we also include a recent implementation of Bayesian nonparametric HMM specifically adapted to step counting. Greedy approaches, such as in [41], rely on exhaustive searches over every possible combination of step numbers and their parameters. Similarly, HMM approaches, such as in [35], rely on filtering procedures that are performed by expensive numerical algorithms. As can be seen in table 2, our method leads not only to dramatically reduced runtimes for every trace tested, but also to better scalability to traces of larger size. Specifically, our results suggest a time-cost that scales almost linearly with trace length, while greedy methods scale almost quadratically. Low runtime is a desirable characteristic for analysis methods, especially in single-molecule assays in which a single experiment typically results in 100-1000 traces that need to be processed individually.

**Table 2:** Computation time comparison between our method, a greed approach [41], and an HMM approach [35]. A baseline trace, similar to the one in fig. 4, was used and its size was then doubled and quadrupled to assess how computational time scales with trace length. Bryan et al.’s method on the longer traces did not complete within the one-hour time limit we allocated.

|  | Computation time |  |  |
| --- | --- | --- | --- |
|  | This study | Kalafut et al. [41] | Bryan et al. [35] |
| Baseline | 13 s | 125 s | 965 s |
| 2x | 28 s | 505 s | - |
| 4x | 51 s | 2164 s | - |

### 3.2 Performance benchmark with data of varying SNR and DER

In our step finding model, the noise, which is represented by the likelihood’s variance, ie eq. (S3) in appendix A.1, is *multiplicative*. That is, the noise on a fluorescent trace varies in tandem with the signal [22, 56, 57]. This characteristic is in stark contrast to additive noise models, where the noise remains uniform over a trace. One reason for the presence of multiplicative noise in our model is the specific operation of cameras and their electronics, which induce nontrivial parameters in the model’s likelihood [23]. However, the largest contribution stems from shot-noise which introduces Poissonian statistics within the model’s likelihood [20, 21].

Due to multiplicative noise, a *single* notion of SNR that precisely characterizes a given fluorescent trace *cannot* be defined. Instead, SNR is specific to each step within a trace. Therefore, to carefully assess the performance of our method across different conditions, we consider the following step-specific definition

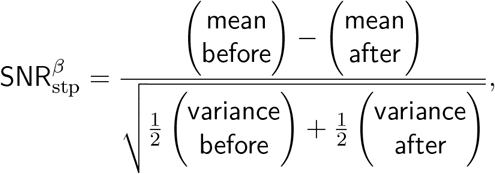

where “before” and “after” indicate the fluorescent measurements over a complete exposure just before and after the time of the *β*^th^ photobleaching event. The precise values for our 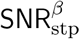 can be computed in terms of the step intensities, background, and the characteristics of the camera and the exposure period; see eq. (S7) for the explicit formula.

To test the robustness of our method in recovering the correct number of steps and also reconstructing the correct signal under different SNR, we generated synthetic data under varying conditions that mimic a wide range of photobleaching experiments. Our synthetic data are obtained by simulating the model of appendix A.1 with prescribed number of steps, background, and step intensities and times. For simplicity, we maintain the same data size, and camera parameters listed in table S1 in agreement with the laboratory data shown in fig. 2. In addition, to facilitate the presentation, we set the intensities of all steps equal to a common value. To mimic different configurations, we generate this value, together with the number of steps, step times, and background *uniformly at random* within the ranges shown in table S4. These ranges include common imaging conditions, such as those that produce the data in fig. 2, and also extreme ones such as those shown in the first and last rows of fig. S3. As expected, our synthetic experiments produce steps with a variety of SNRs that range from ≈1 to ≈8. As a reference, the SNRs of the steps in fig. 2 range from ≈4 to ≈8.

We analyze each synthetic trace using our method and quantify the results using two key *metrics*.

First, the absolute difference

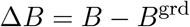

in the total number *B* of steps recovered from its respective ground truth *B*^grd^. Second, the relative difference

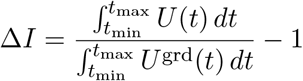

in the total integration of the signal *U* (*t*) recovered from its respective ground truth *U* ^grd^(*t*). For these metrics, *B* and *U* (*t*) are obtained via eq. (S5) and eq. (S4), respectively. Our first metric focuses exclusively on step counting, but does not quantify the ability of our method to accurately estimate the correct signal characteristics such as step intensities, times, and background. Our second metric is more restrictive and accurately captures deviations in photon emission rates and step times in addition to the total step count.

Figure 10 demonstrates both metrics obtained from our synthetically generated traces. For clarity, we summarize each synthetic trace by its minimum SNR^min^ and maximum SNR^max^. As can be seen, our method remains robust across a wide range of SNR. Specifically, for SNR’s greater than ≈2 both metrics are tightly concentrated around 0 indicating that the number of recovered steps or the reconstructed signal agree with their ground truths with great confidence. Our method yields acceptable results even for SNR below 2 (for characteristic traces see upper row of fig. S3); however, these cases are associated with larger uncertainty and occasional under- or over-estimation of steps.

**Figure 10.**
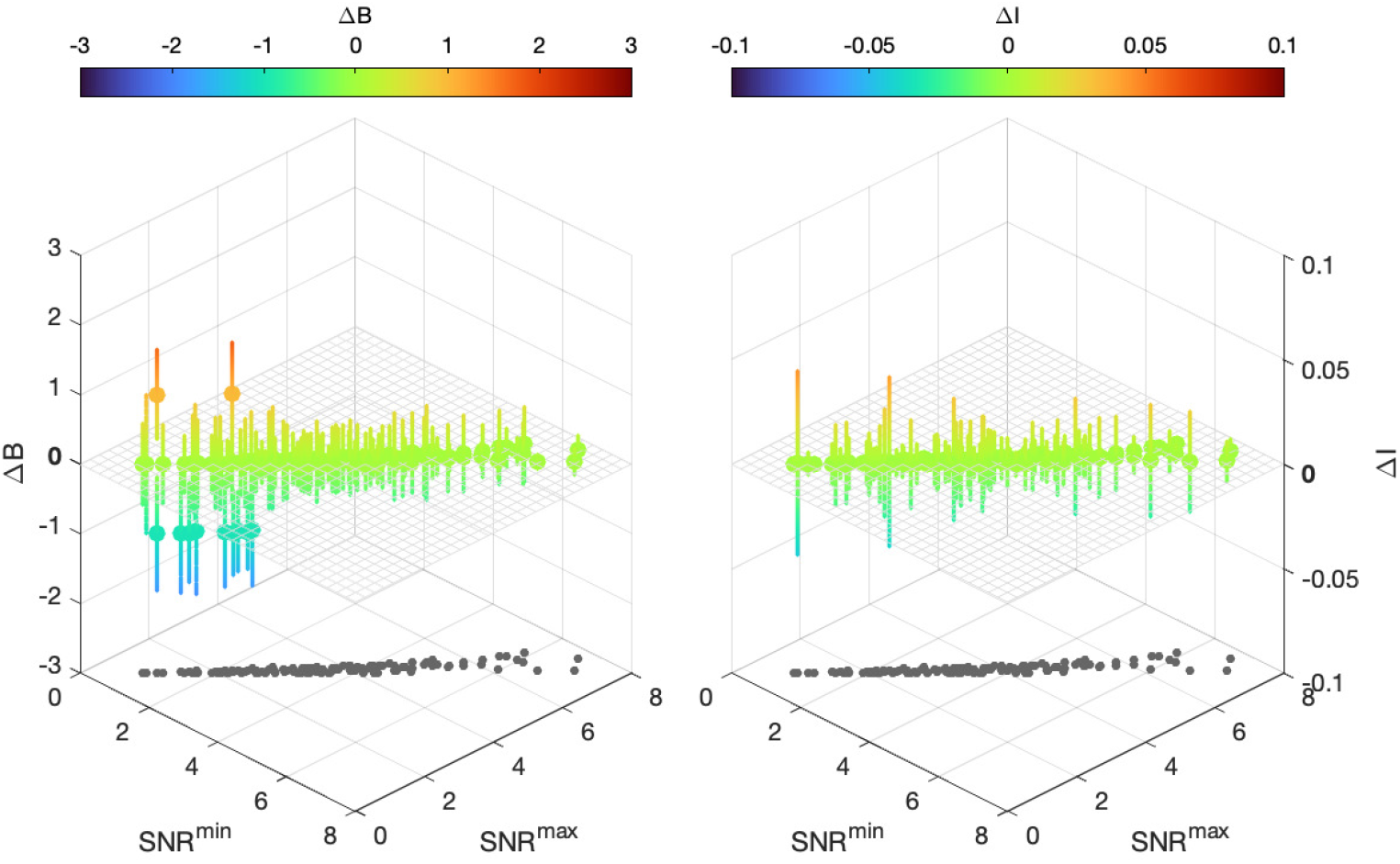
Sampler performance across data of varying SNR for 150 random experiments, with experimental conditions sampled uniformly from the ranges specified in table S4. On the left panel we show the median absolute difference between the recovered and ground truth total number of step (ΔB); while on the right, we show the median of ΔI as defined in the main text. Error bars represent one standard deviation.

Besides SNR, another critical indicator of the quality of a fluorescent trace is the duration of the time period that individual steps are visible. To accurately quantify such an indicator, we consider a duration-to-exposure metric. For each step, this is given by

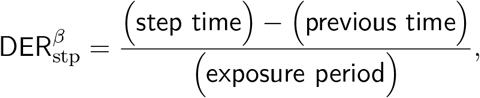

where “step time” and “previous time” denote the time of the *β*^th^ and its predecessor photobleaching events, respectively. In this definition, we normalize with respect to the exposure period, so we obtain a unitless number that better correlates with the number of data points related to each event. The precise values for our 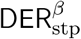 can be computed in terms of the timing schedule of the camera acquiring the measurements and the exposure period; see eq. (S8) for the explicit formula.

Figure 11 shows how our step estimation Δ*B* and signal reconstruction Δ*I* metrics of our synthetically generated traces correlate with DER. Similarly to the preceding presentation, for clarity, we summarize each synthetic trace by its minimum DER^min^ and maximum DER^max^. As can be seen, our method remains robust across a wide range of DER. Specifically, for DER’s greater than ≈ 10 both metrics are concentrated around 0 indicating that the number of recovered steps or the reconstructed signal agree with their ground truths with great confidence. Our method yields acceptable results even for DER below 10; however, these cases are associated with an occasional underestimation of steps, especially for those traces that suffer from low SNR.

**Figure 11.**
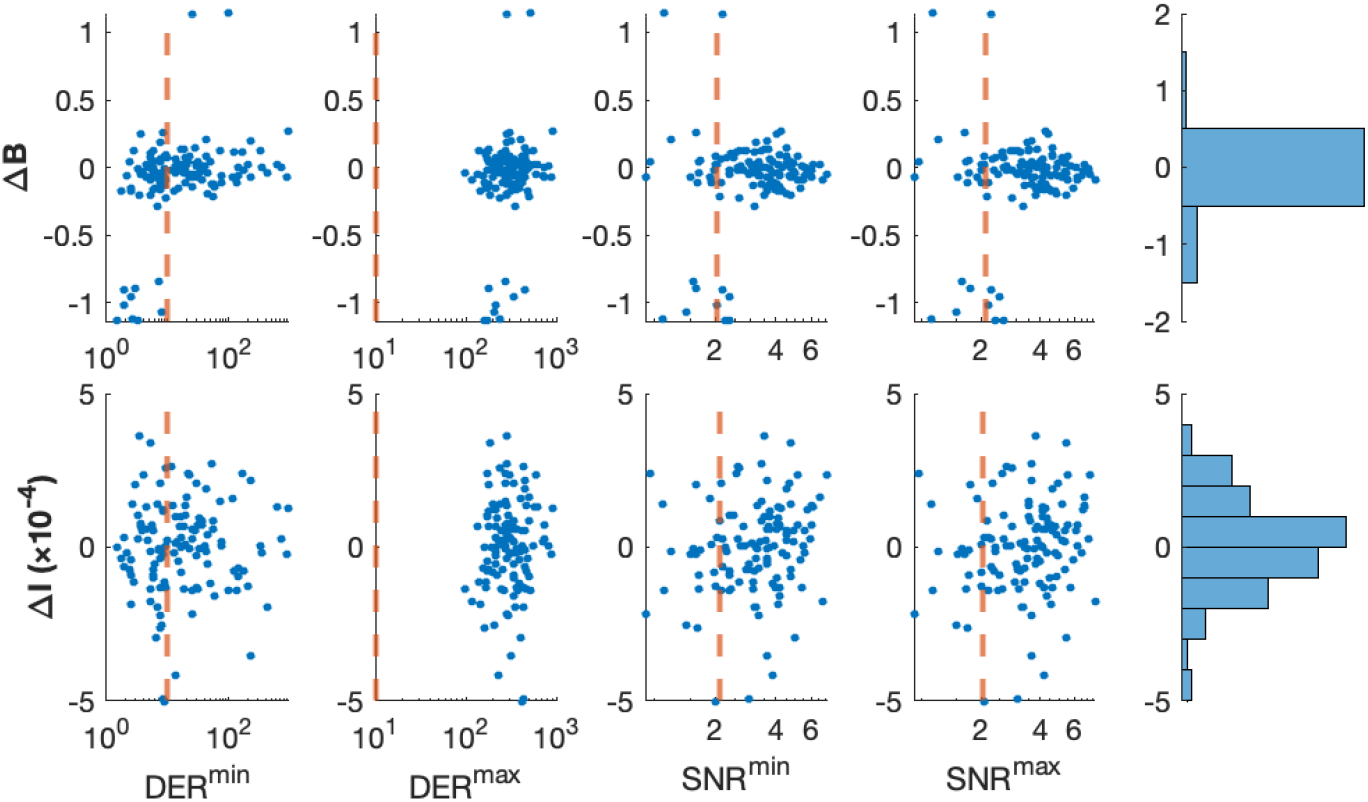
Relationship between step estimation error ΔB (top row) and signal reconstruction error ΔI (bottom row) with data quality metrics: DER^min^, DER^max^, SNR^min^, and SNR^max^. Vertical lines, at 10 for DER and 2 for SNR, mark reference thresholds. For visualization purposes, small jitter is added to ΔB values and quality metrics are plotted on a logarithmic scale.

## 4 Discussion

Analysis of single-molecule fluorescence traces with an efficient statistical framework is an essential requirement to determine the biological functions and dynamics of molecular clusters within diffraction-limited regions [19, 24, 29]. Following a photobleaching experiment, this task reduces to accurately and quickly determining the number of steps present in the acquired fluorescence time series. This allows investigators to deduce the stoichiometry of the molecules under study and gain insight into their structural and functional properties. The work we present here provides a novel statistical model to represent fluorescence traces and four different algorithmic procedures to analyze them. Our methods not only determine the total number of steps, but also provide full posterior distributions for individual steps and their characteristics. We do so without relying on restrictive assumptions on the dynamics of the molecules or fluorophores under study, instead providing a kinetics-independent approach that can be extended to a variety of single-molecule fluorescence experiments.

Our methods were illustrated by applying them to raw data from TIRF microscopy experiments. We demonstrated our method’s ability to recover the number of steps present and validated our method by reproducing the experimental settings using synthetically generated data. We showed that our method is able to accurately recover ground truth in both low- and high-SNR scenarios. Moreover, we evaluated the quality of our methods by evaluating their mixing properties and found that our sampler is able to efficiently explore the parameter space, converges rapidly, and has low autocorrelation. Furthermore, we challenged our methods by conducting extensive benchmarking tests covering a wide spectrum of photobleaching conditions and demonstrating that it can serve as a general purpose tool across SNR. Finally, a head-to-head comparison with existing methodologies demonstrated its advantages in leading to artifact-free unbiased estimates, and its superior computational performance that enables large-scale analysis efforts.

The contribution that this work brings will particularly apply to data sets that are effected by higher degrees of noise and have a weaker signature, allowing practitioners to tackle experiments that have been inaccessible before. Data acquired at the single-molecule level, such as the SiMPull-POP data explored here, are often accompanied by high background and noise, which can obscure and convolute the data analysis process. By applying a fully automated statistical approach that is independent of user’s subjective input, as we have shown here, it can both confirm and compare current data quantification methods to validate experimental data results. In addition, it will help reduce potential data quantification bias, reveal photobleaching steps that are difficult to distinguish by visual inspection, and provide a more detailed characterization of the data at hand.

Although in this study we applied our sampler to a specific experimental data set, the methodology we propose can be generalized and applied more broadly within the sphere of photobleaching step analysis. However, despite its novelty and computational efficiency, our method presents certain limitations. First, it is mathematically demanding, so its adoption is based on existing software. We provide a prototype implementation (source code and graphical user interface) in Supporting Materials and in our GitHub repository [58]. Second, our method is sensitive to experimental details related to data acquisition hardware. Currently, our implementation supports fluorescence data from one-channel recordings acquired via wide-field microscopes equipped with either EMCCD or CMOS cameras. Nevertheless, two-color imaging, FRET, or even rapid single-photon recordings can be incorporated, but only with nontrivial modifications to our detection setup. Obviously, such cases require problem-specific implementations that will be the focus of future studies.

## 5 Methods

### 5.1 Data analysis

We develop a specialized statistical model for data analysis that includes physically faithful signal, stimulus, and noise representations. The model that we formulate does *not* use kinetic schemes or other dynamical representations, so it can be applied in situations where the dynamics probed may or may not be Markovian. We apply Bayesian principles to provide parameter estimates and provide specialized computational procedures in order to evaluate the values of our estimators. That is, we develop four different MCMC samplers with increasing sophistication. For details, see formulations in Supplement. Specifically, refer to appendices A.1 and A.3 for our model formulation, and appendices A.5 and B for our computational approaches.

### 5.2 Data acquisition

For the demonstrations shown in Results, we used synthetic and real-life data from single-molecule experiments.

The *laboratory data* are obtained by the SiMPull-POP TIRF photobleaching assay, as detailed in [19] of a full-length EphA2 fused to the C-terminal GFP that was stably expressed in mammalian cells DU145 (ATCC^®^ HTB-81).

The *synthetic data* is obtained by simulating the model of appendix A.1 using standard pseudorandom operations. For the simulations, we set the timing and camera parameters according to laboratory settings (see above) and generated measurements of equal size spanning an equivalent period of real time. We list our specific values in appendix E. For all simulations, we prescribe the values of *c*_bck_, *B*, 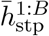, and 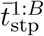. The ground truth values are maintained for comparison with MCMC, but otherwise not used.

## Acknowledgments

Lamichhane laboratory research is supported by the National Institutes of Health R35GM142946 (R.L). Barrera laboratory research is supported by the National Institutes of Health R35GM140846 (F.N.B).

## Author contributions

C.M. contributed to the analysis methods, computational implementation and software development.

C.M. and I.S. prepared the manuscript. A.W., S.T.K., R.L., and F.N.B. provided the experimental data and commented on the manuscript. I.S. conceived the research and supervised all aspects of the project.

