## Supplement for "Bayesian analysis and efficient algorithms for single-molecule fluorescence data and step counting"

**C Measurements and signal reduction** **12**

**D Data quality metrics** **12**

**E Parameters, hyperparameters, and other settings** **13**

**F Sensitivity analysis** **15**

**G Additional Figures** **15**

### A Methods of data analysis

Here, we describe the formulation of our model formulation for the representation of raw photobleaching step data, present our Bayesian framework for its analysis, and outline the computational approaches that we apply for their evaluation.

#### A.1 Model formulation

We consider the analysis of raw photobleaching step data that are acquired in a single experiment session. Our experiment begins at time $t_{\min}$ and ends at time $t_{\max}$ during which photodetection measurements $w_n$ are collected at respective times $t_n$, which we index by *subscripts* $n = 1, \ldots, N$. Most often, the acquisition times are regularly spaced as determined by the exposure period $\tau_{\text{exp}}$ and the dead period $\tau_{\text{dead}}$ adopted in the experiment, e.g. $t_n = t_{n-1} + \tau_{\text{exp}} + \tau_{\text{dead}}$. Although we do not necessarily require such regularly spaced times in our analysis, by convention, we require that these be strictly ordered so that $t_{\min} \leq t_1 < t_2 < \cdots < t_N \leq t_{\max}$. In this way, our raw measurements take the form of two sorted time series $t_{1:N}$ and $w_{1:N}$.

Similarly to the existing step finding techniques [11,12], we employ a detection model that considers the signal, stimulus, and noise components of each measurement. The signal model represents steps in continuous time and is given by

$$U(t) = c_{\text{bck}} + \sum_{\beta=1}^{B} h_{\text{stp}}^{\beta} H(t - t_{\text{stp}}^{\beta}), \tag{S1}$$

where $c_{\text{bck}}$ is the background signal which we assume remains constant throughout the experiment, $B$ is the total number of steps, $h_{\text{stp}}^{\beta}$ is the decrease in signal following the $\beta^{\text{th}}$ step, and $t_{\text{stp}}^{\beta}$ is the time at which the $\beta^{\text{th}}$ step occurs. The function $H(t)$ models the step response. Specifically, we use $H(t) = 1$ for $t < 0$ to model fluorescence and $H(t) = 0$ for $t > 0$ to model photobleaching. In this way, each summand $h_{\text{stp}}^{\beta} H(t - t_{\text{stp}}^{\beta})$ in eq. (S1) represents an individual photobleaching event that occurs at time $t_{\text{stp}}^{\beta}$ of intensity $h_{\text{stp}}^{\beta}$.

We index the steps with *superscripts* $\beta = 1, \ldots, B$ and, in order to create a unique labeling among the $B!$ possible permutations of our model steps, we consider only signals with distinct time steps and labeling from early to late events such that

$$t_{\text{stp}}^{1} < t_{\text{stp}}^{2} < \cdots < t_{\text{stp}}^{B}. \tag{S2}$$

To link our signal with measurements acquired in a typical photobleaching experiment, we model integrative detection with the stimulus model proposed in recent HMJP studies [11, 12]. Hence, for each measurement $w_n$, we model an associated stimulus $u_n$ determined by

$$u_n = \int_{t_n - \tau_{\text{exp}}/2}^{t_n + \tau_{\text{exp}}/2} U(t)\, dt,$$

where $\tau_{\text{exp}}$ is the exposure period which accounts for the amount of time the detector acquiring each measurement remains responsive to incident light. Lastly, we assume multiplicative Gaussian noise for the measurements $w_n$ and let

$$w_n | u_n \sim \text{Normal}\left(\mu + g u_n, v + f g^2 u_n\right), \tag{S3}$$

where $\mu, v, g$ and $f$ are camera specific parameters, namely the read-out offset, read-out variance, overall gain, and excess noise factor respectively. As explained in [31], this is a generic detection model that accounts for fluorescence measurements contaminated by shot noise and readout noise obtained with EMCCD or CMOS cameras. The readout parameters $\mu, v, g, f$ can be obtained by following the protocols in [31] and combining the response of all pixels whose intensity is summed up, yielding the measurements $w_{1:N}$.

#### A.2 Model reparameterization

Because the total number of steps $B$ in our signal is an unknown quantity of interest, eq. (S1) requires reformulation. For this, similar to recent non-parametric studies [2, 4, 13, 34], we begin by considering a total of $M$ model steps and, for each model step, introduce an indicator variable $\bar{b}^m_{\text{stp}}$ that may only take values $0$ or $1$. Steps whose corresponding indicator variable is $1$ are considered active, while those with indicator variable $0$ are considered inactive.

Introducing these indicator variables allows us to consider an arbitrarily large number of model steps, with $M \gg B$, and estimate the number of active steps simultaneously with the rest of the parameters by treating each $\bar{b}^m_{\text{stp}}$ as a separate parameter and estimating its value. With this modification, eq. (S1), becomes,

$$U(t) = c_{\text{bck}} + \sum_{m=1}^{M} \bar{b}^m_{\text{stp}} \bar{h}^m_{\text{stp}} H(t - \bar{t}^m_{\text{stp}}). \tag{S4}$$

This way, the total number of steps $B$ is now a variable, which we obtain by

$$B = \sum_{m=1}^{M} \bar{b}^m_{\text{stp}}, \tag{S5}$$

and can be included in our Bayesian nonparametric setup (described below).

Additionally, due to the summation in eq. (S1), any permutation of the $\beta$ indices leads to the same signal $U(t)$, making the model non-identifiable *without* the conditions in eq. (S2), [19]. These conditions negatively affect the performance of the Markov Chain Monte Carlo (MCMC) samplers [3, 5, 8]. Hence, in preparation for the computational implementation of our model (see appendix A.5), we drop the constraints in eq. (S2) and introduce a different indexing convention in eq. (S4).

Converting the indexing from $\beta$ to $m$ improves the mixing of the MCMC samplers (described next); however, due to the nonidentifiability of the model, we need additional post-processing to recover the original labeling. Specifically, starting from the vector $\bar{t}^m_{\text{stp}}$, we select the entries that correspond to active steps and reorder them in ascending order to ensure that they follow the order in which the steps occur. These become the entries of vector $t^\beta_{\text{stp}}$. Then, we let $h^{1:B}_{\text{stp}}$ be the steps intensities selected from $\bar{h}^m_{\text{stp}}$ that correspond to times $t^{1:B}_{\text{stp}}$. This creates a relabeling map $\left(\bar{b}^{1:M}_{\text{stp}}, \bar{t}^{1:M}_{\text{stp}}, \bar{h}^{1:M}_{\text{stp}}\right) \mapsto \left(B, t^{1:B}_{\text{stp}}, h^{1:B}_{\text{stp}}\right)$ that abides by the constraint in eq. (S2).

#### A.3 Bayesian considerations

The unknown quantities of interest in our model are $c_{\text{bck}}, \bar{h}^{1:M}_{\text{stp}}, \bar{t}^{1:M}_{\text{stp}}$, and $\bar{b}^{1:M}_{\text{stp}}$. To allow estimation of their values from raw measurements, we follow the Bayesian principles [6, 18] and apply the priors that follow:

$$\begin{aligned}
c_{\text{bck}} &\sim \text{Gamma}\left(\phi_{\text{bck}}, c^{\text{ref}}_{\text{bck}}/\phi_{\text{bck}}\right), && \\
\bar{h}^m_{\text{stp}} &\sim \text{InvGamma}\left(\phi_{\text{stp}}, (\phi_{\text{stp}} - 1)h^{\text{ref}}_{\text{stp}}\right), && m = 1, \ldots, M, \\
\bar{t}^m_{\text{stp}} &\sim \text{Uniform}_{[t^{\min}_{\text{stp}}, t^{\max}_{\text{stp}}]}, && m = 1, \ldots, M, \\
\bar{b}^m_{\text{stp}} &\sim \text{Bernoulli}(\gamma/M), && m = 1, \ldots, M.
\end{aligned}$$

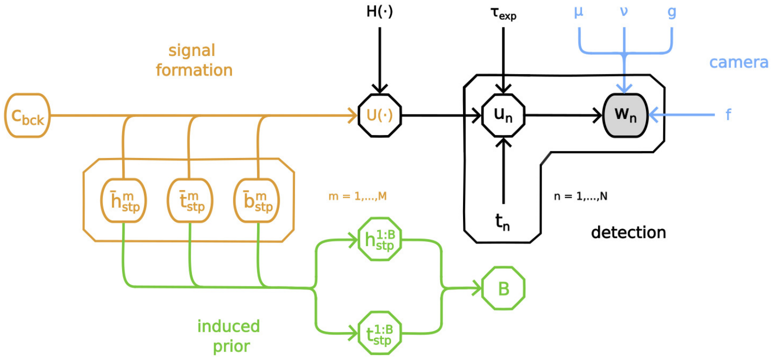

*Figure S1: A graphical model illustrating the random variables and parameters used in the statistical analysis of photobleaching data. The main detection model is shown in black, the camera-specific parameters are shown in blue, the signal formation portion of the model is in orange, and the induced prior for the total active steps variable $B$ is shown in green.*

Because after reparameterization the total number of steps $B$ becomes a derived quantity of interest, similar to other nonparametric models [1, 9, 10, 22–30, 33, 34], it receives an induced prior. As the sum of independent and identically distributed Bernoulli trials [18], this is given by

$$B \sim \text{Binomial}(M, \gamma/M).$$

According to the law of rare events [18], this distribution converges to Poisson($\gamma$) as $M \to \infty$ which, similarly to other Bayesian nonparametric statistical methods, indicates that our model avoids overfitting at $M \gg 1$.

A graphical summary of the full Bayesian model that we implement, including signal formation and applied and induced priors, is shown in fig. S1.

#### A.4 Summary of model equations

Our entire statistical model consists of the following equations.

$$\begin{aligned}
c_{\text{bck}} &\sim \text{Gamma}\left(\phi_{\text{bck}}, c^{\text{ref}}/\phi_{\text{bck}}\right), \\
\bar{h}^m_{\text{stp}} &\sim \text{InvGamma}\left(\phi_{\text{stp}}, (\phi_{\text{stp}}-1)h^{\text{ref}}_{\text{stp}}\right), && m = 1,\dots,M, \\
\bar{t}^m_{\text{stp}} &\sim \text{Uniform}_{[t^{\min}_{\text{stp}}, t^{\max}_{\text{stp}}]}, && m = 1,\dots,M, \\
\bar{b}^m_{\text{stp}} &\sim \text{Bernoulli}\left(\gamma/M\right), && m = 1,\dots,M, \\
w_n | c_{\text{bck}}, \bar{h}^{1:M}_{\text{stp}}, \bar{t}^{1:M}_{\text{stp}}, \bar{b}^{1:M}_{\text{stp}} &\sim \text{Normal}\left(\mu + g u_n, v + f g^2 u_n\right), && n = 1,\dots,N.
\end{aligned}$$

In these equations, the stimuli, effective step responses, and step response are obtained according to

$$u_n = \tau_{\text{exp}} c_{\text{bck}} + \sum_{m=1}^{M} \bar{b}^m_{\text{stp}} \bar{h}^m_{\text{stp}} G_n\left(\bar{t}^m_{\text{stp}}\right), \quad G_n(t_{\text{stp}}) = \int_{t_n-\tau_{\text{exp}}/2}^{t_n+\tau_{\text{exp}}/2} H(t-t_{\text{stp}})dt, \quad H(t) = \begin{cases} 1, & t < 0. \\ 0, & t > 0. \end{cases}$$

#### A.5 Computational approaches

The posterior distribution of our model $p(c_{\text{bck}}, h^{1:B}_{\text{stp}}, t^{1:B}_{\text{stp}} | w_{1:N})$, which we use to derive estimates [6, 18], cannot be analytically characterized. Hence, we use Monte Carlo methods [18, 21].

#### A.5.1 Overview

Our Monte Carlo scheme proceeds by first completing the targeted distribution with respect to the characteristics of the model step $\bar{h}^{1:M}_{\text{stp}}, \bar{t}^{1:M}_{\text{stp}}, \bar{b}^{1:M}_{\text{stp}}$. Formally, our completion is as follows

$$p(c_{\text{bck}}, h^{1:B}_{\text{stp}}, t^{1:B}_{\text{stp}}|w_{1:N}) = \int \cdots \int p(c_{\text{bck}}, h^{1:B}_{\text{stp}}, t^{1:B}_{\text{stp}}|\bar{h}^{1:M}_{\text{stp}}, \bar{t}^{1:M}_{\text{stp}}, \bar{b}^{1:M}_{\text{stp}})p(c_{\text{bck}}, \bar{h}^{1:M}_{\text{stp}}, \bar{t}^{1:M}_{\text{stp}}, \bar{b}^{1:M}_{\text{stp}}|w_{1:N}),$$

where integration is performed over the values of $\bar{h}^{1:M}_{\text{stp}}, \bar{t}^{1:M}_{\text{stp}}, \bar{b}^{1:M}_{\text{stp}}$. We then, on the completed target, apply two-stage ancestral sampling [18]. First, we sample $p(c_{\text{bck}}, \bar{h}^{1:M}_{\text{stp}}, \bar{t}^{1:M}_{\text{stp}}, \bar{b}^{1:M}_{\text{stp}}|w_{1:N})$ using MCMC methods, and then sample $p(c_{\text{bck}}, h^{1:B}_{\text{stp}}, t^{1:B}_{\text{stp}}|\bar{h}^{1:M}_{\text{stp}}, \bar{t}^{1:M}_{\text{stp}}, \bar{b}^{1:M}_{\text{stp}})$ directly through the relabeler of appendix A.2.

For the MCMC stage, which is the computationally most demanding, we develop *four variations* of a Gibbs sampler [18, 21]. As discussed in the following, we base these samplers on a naive approach and introduce modifications of increasing complexity and sophistication in successive layers.

The most rudimentary version of our sampler updates one variable at a time using a standard Metropolis-Hastings (MH) algorithm with additive random walks [21]. Due to the shortcomings of this approach, as discussed in more detail in Results, we introduce an initial set of modifications based on the existing literature. For example, we replace the MH algorithm to sample $\bar{t}^{1:M}_{\text{stp}}$ with a slice sampler [16] and opt for multiplicative random walks [21] in the MH schemes for $\bar{h}^{1:M}_{\text{stp}}$ and $c_{\text{bck}}$. However, despite some improvements, this sampler still leads to inadequate performance. With our third sampler, we update multiple parameters simultaneously. In particular, we perform a stochastic update on one variable and a simultaneous deterministic update on another [18], with the aim of directing the sampler towards areas of high probability. Although this sampler appears adequate for practical applications, its performance is critically dependent on manual fine-tuning of the involved MH steps. Finally, in our most advanced version of the sampler, we eliminate any reliance on the tuning of the proposal parameters by replacing the MH joint updates for $c_{\text{bck}}$ and $\bar{h}^{1:M}_{\text{stp}}$ with elliptical multiplicative slice samplers. Next, we describe our methods in more detail.

#### A.5.2 Gibbs sampling schemes

Each of our Gibbs sampling schemes targets $p(c_{\text{bck}}, \bar{h}^{1:M}_{\text{stp}}, \bar{t}^{1:M}_{\text{stp}}, \bar{b}^{1:M}_{\text{stp}}|w_{1:N})$ and consists of two basic stages:

- Initialize $c_{\text{bck}}, \bar{h}^{1:M}_{\text{stp}}, \bar{t}^{1:M}_{\text{stp}}$, and $\bar{b}^{1:M}_{\text{stp}}$ by randomly sampling their respective priors.
- Iterate the steps described next in random order until a sufficient number of samples are produced, storing samples after each iteration.

A summary of the iteration stage of each scheme is provided in the following, and a detailed breakdown of each sampler, together with the necessary implementation details, is provided in appendix B.

**Sampler 1**

- Update $c_{\text{bck}}$ by sampling its full conditional using an *additive Metropolis* random walk.
- Update each one of $\bar{h}^{1:M}_{\text{stp}}$, in random order, by sampling its full conditional using an *additive Metropolis* random walk.
- Update each one of $\bar{t}^{1:M}_{\text{stp}}$, in random order, by sampling its full conditional using an *additive Metropolis* random walk.
- Update each one of $\bar{b}^{1:M}_{\text{stp}}$, in random order, by *sampling directly* its full conditional.

**Sampler 2**

- Update $c_{\text{bck}}$ by sampling its full conditional using a *multiplicative Metropolis* random walk.

- Update each one of $\bar{h}^{1:M}_{\rm stp}$, in random order, by sampling its full conditional using a *multiplicative Metropolis* random walk.
- Update each one of $\bar{t}^{1:M}_{\rm stp}$, in random order, by sampling its full conditional using a *slice sampler.*
- Update each one of $\bar{b}^{1:M}_{\rm stp}$, in random order, by *sampling directly* its full conditional.

**Sampler 3**

- Update $c_{\rm bck}$ by sampling its full conditional using a *multiplicative Metropolis* random walk and simultaneously make a *deterministic adjustment* in the last active step, if there is one.
- Update each one of $\bar{h}^{1:M}_{\rm stp}$, in random order, by sampling its full conditional using a *multiplicative Metropolis* random walk and simultaneously making a *deterministic adjustment* in the previous active step, if there is one.
- Update each one of $\bar{t}^{1:M}_{\rm stp}$, in random order, by sampling its full conditional using a *slice sampler.*
- Update each one of $\bar{b}^{1:M}_{\rm stp}$, in random order, by *sampling directly* its full conditional and simultaneously make a *deterministic adjustment* on either the previous or the following active step, if there is one. If there is no following active step, make a deterministic adjustment on the background.

**Sampler 4**

- Update $c_{\rm bck}$ by sampling its full conditional using a *multiplicative slice sampler* and simultaneously making a *deterministic adjustment* in the last active step, if there is one.
- Update each one of $\bar{h}^{1:M}_{\rm stp}$, in random order, by sampling its full conditional using a *multiplicative slice sampler* random walk and simultaneously making a *deterministic adjustment* in the previous active step, if there is one.
- Update each one of $\bar{t}^{1:M}_{\rm stp}$, in random order, by sampling its full conditional using a *slice sampler.*
- Update each one of $\bar{b}^{1:M}_{\rm stp}$, in random order, by *sampling directly* its full conditional and simultaneously make a *deterministic adjustment* on either the previous or the following active step, if there is one. If there is no following active step, make a deterministic adjustment on the background.

### B Implementation of individual samplers

To improve readability, we simplify our notation in this section only and denote our variables $c_{\rm bck}$, $\bar{h}^m_{\rm stp}$, $\bar{t}^m_{\rm stp}$, $\bar{b}^m_{\rm stp}$ with $c, \bar{h}^m, \bar{t}^m, \bar{b}^m$.

#### B.1 Sampler 1: A naive Gibbs scheme

Our first sampler consists of basic textbook MCMC sampling procedures. Improvements to this scheme are gradually introduced and discussed in the following sections.

##### B.1.1 Sampling the background

We aim to sample $c$ from its full conditional $p(c|\bar{t}^{1:M}, \bar{h}^{1:M}, \bar{b}^{1:M}, w_{1:N})$. For this, we use an MH sampler. Our proposals are additive random walks of the form

$$c^{\rm prop}|c^{\rm old} \sim \text{Normal}\left(c^{\rm old}, \sigma_c^2\right),$$

where $\sigma_c$ is an adjustable parameter. We tune this parameter during burn-in [20] so that the resulting acceptance rate falls within $25\%$ to $50\%$.

##### B.1.2 Sampling the step intensities

We aim to sample $\bar{h}^m$, where $m$ is chosen uniformly at random among $1, \ldots, M$, from its full conditional $p(\bar{h}^m|c, \bar{t}^{1:M}, h^{-m}, \bar{b}^{1:M}, w_{1:N})$. Depending on $\bar{b}^m$, we apply one of the choices:

- If $\bar{b}^m = 0$, our full conditional reduces to the prior $p(\bar{h}^m)$ which we sample directly.
- If $\bar{b}^m = 1$, we use an MH sampler. Our proposals are additive random walks of the form

$$(\bar{h}^m)^{\text{prop}} | (\bar{h}^m)^{\text{old}} \sim \text{Normal}\left((\bar{h}^m)^{\text{old}}, \sigma_h^2\right),$$

where $\sigma_h$ is an adjustable parameter. We tune this parameter during burn-in [20] so that the resulting acceptance rate falls within $25\%$ to $50\%$.

##### B.1.3 Sampling the step times

We aim to sample $\bar{t}^m$, where $m$ is chosen uniformly at random among $1, \ldots, M$, from its full conditional $p(\bar{t}^m | c, \bar{h}^{1:M}, \bar{t}^{-m}, \bar{b}^{1:M}, w_{1:N})$. Depending on $\bar{b}^m$, we apply one of the choices:

- If $\bar{b}^m = 0$, our full conditional reduces to the prior $p(\bar{t}^m)$ which we sample directly.
- If $\bar{b}^m = 1$, we use an MH sampler. Our proposals are additive random walks of the form

$$(\bar{t}^m)^{\text{prop}} | (\bar{t}^m)^{\text{old}} \sim \text{Normal}\left((\bar{t}^m)^{\text{old}}, \sigma_t^2\right),$$

where $\sigma_t$ is an adjustable parameter. We tune this parameter during burn-in [20] so that the resulting acceptance rate falls within $25\%$ to $50\%$.

##### B.1.4 Sampling the step loads

We aim to sample $\bar{b}^m$, where $m$ is chosen uniformly at random among $1, \ldots, M$, from its full conditional $p(\bar{b}^m | c, \bar{h}^{1:M}, \bar{t}^{1:M}, \bar{b}^{-m}, w_{1:N})$ using an MH sampler. Our proposals are of the form

$$(\bar{b}^m)^{\text{prop}} | (\bar{b}^m)^{\text{old}} \sim \text{Bernoulli}\left(1 - (\bar{b}^m)^{\text{old}}\right).$$

We note that our approach is equivalent to direct sampling [18]; however, we opt for an MH format in order to facilitate the presentation of the samplers that follow.

#### B.2 Sampler 2: An improved Gibbs scheme

Here we formulate an improved Gibbs sampler which proceeds similarly to the naive one of appendix B.1.

##### B.2.1 Sampling the background

We proceed as in appendix B.1.1; however, we replace the additive random-walk proposals with symmetric multiplicative ones. As described in [18], these are obtained by

$$\begin{aligned}\epsilon^{\text{prop}} &\sim \frac{1}{2}\text{Beta}(\alpha_c, 1) + \frac{1}{2}\text{InvBeta}(\alpha_c, 1),\\ c^{\text{prop}} &= c^{\text{old}}\epsilon^{\text{prop}},\end{aligned}$$

where $\alpha_c$ is an adjustable parameter. We tune this parameter during burn-in [20] so that the resulting acceptance rate falls within $25\%$ to $50\%$.

##### B.2.2 Sampling the step intensities

We proceed as in appendix B.1.2; however, we replace the additive random-walk proposals with symmetric multiplicative ones. As described in [18], these are obtained by

$$\begin{aligned}\epsilon^{\text{prop}} &\sim \frac{1}{2}\text{Beta}(\alpha_h, 1) + \frac{1}{2}\text{InvBeta}(\alpha_h, 1),\\ (\bar{h}^m)^{\text{prop}} &= (\bar{h}^m)\epsilon^{\text{prop}},\end{aligned}$$

where $\alpha_h$ is an adjustable parameter. We tune this parameter during burn-in [20] so that the resulting acceptance rate falls within $25\%$ to $50\%$.

#### B.2.3 Sampling the step times

We proceed as in appendix B.1.3; however, for the case $\bar{b}^m = 1$, we replace the additive random-walk proposals with a fixed-interval slice sampler as described in [16]. Our initial interval is determined by the prior's hyperparameters $t_{\mathrm{stp}}^{\mathrm{min}}$ and $t_{\mathrm{stp}}^{\mathrm{max}}$.

#### B.2.4 Sampling the step loads

We proceed as in appendix B.1.4.

### B.3 Sampler 3: An advanced Gibbs scheme

In the samplers of appendices B.1 and B.2, only one variable is updated at a time. As shown in RESULTS, this leads to practical non-ergodicity, where the sampler has difficulty exploring the target distribution efficiently and gets trapped in local maxima. Hence, we propose a *novel sampler* that is able to explore the parameter space by updating multiple, carefully selected, variables jointly. Moreover, we guide the sampler towards areas of high probability by utilizing transformations of the selected variables that correspond to approximately the same stimuli over large signal periods and, therefore, have high acceptance. To obtain such adjustments, in general, we update variables in pairs with one stochastic and one deterministic proposal.

Before we describe the fine details of our sampling procedures, we present an overview of our novel sampling mechanism. For this, we start with an MH procedure on a generic target $\pi(x, y)$ for a pair of scalar variables $x, y$. Our algorithm is based on a family of proposal mechanisms, parametrized by some $\epsilon > 0$, of the form:

$$x^{\mathrm{prop}} | x^{\mathrm{old}} \sim V_{x^{\mathrm{old}}},$$
$$y^{\mathrm{prop}} | x^{\mathrm{prop}}, x^{\mathrm{old}}, y^{\mathrm{old}} \sim \mathsf{Normal}(y^{\mathrm{old}} + f(x^{\mathrm{prop}}, x^{\mathrm{old}}), \epsilon).$$

With this scheme, our aim is to propose $x$ through a distribution $V_{x^{\mathrm{old}}}$ as usual and adjust $y$ in response to $x$ through an appropriate function $f(\cdot, \cdot)$ to increase the acceptance of the proposals and thus improve the search strategy over the parameter space.

For any member of this family, the MH acceptance ratio [18, 20, 21] is given by

$$\alpha_\epsilon = \frac{\pi(x^{\mathrm{prop}}, y^{\mathrm{prop}})}{\pi(x^{\mathrm{old}}, y^{\mathrm{old}})} \frac{V_{x^{\mathrm{prop}}}(x^{\mathrm{old}})}{V_{x^{\mathrm{old}}}(x^{\mathrm{prop}})} \frac{\mathsf{Normal}(y^{\mathrm{old}}; y^{\mathrm{prop}}) + f(x^{\mathrm{old}}, x^{\mathrm{prop}}), \epsilon)}{\mathsf{Normal}(y^{\mathrm{prop}}; y^{\mathrm{old}} + f(x^{\mathrm{prop}}, x^{\mathrm{old}}), \epsilon)}.$$

From now on, we *only* consider adjustment functions $f(\cdot, \cdot)$ that are skew-symmetric, meaning

$$f(x', x'') = -f(x'', x').$$

In this special case, the last factor in $\alpha_\epsilon$ reduces to $1$, and so the acceptance ratio simplifies to

$$\alpha_\epsilon = \frac{\pi(x^{\mathrm{prop}}, y^{\mathrm{prop}})}{\pi(x^{\mathrm{old}}, y^{\mathrm{old}})} \frac{V_{x^{\mathrm{prop}}}(x^{\mathrm{old}})}{V_{x^{\mathrm{old}}}(x^{\mathrm{prop}})}. \tag{S6}$$

Since $\alpha_\epsilon$ is now independent of $\epsilon$, the same ratio carries over even at the *limit* $\epsilon \to 0$. At the same limit, our proposal mechanism for $y^{\mathrm{prop}}$ degenerates, as desired, to a deterministic choice

$$y^{\mathrm{prop}} = y^{\mathrm{old}} + f(x^{\mathrm{prop}}, x^{\mathrm{old}}).$$

In summary, we have shown that, provided that $f(\cdot, \cdot)$ is skew-symmetric, our proposal mechanism

$$x^{\mathrm{prop}} | x^{\mathrm{old}} \sim V_{x^{\mathrm{old}}},$$
$$y^{\mathrm{prop}} = y^{\mathrm{old}} + f(x^{\mathrm{prop}}, x^{\mathrm{old}}).$$

results in a valid MH sampler and its acceptance ratio is given by eq. (S6).

#### B.3.1 Sampling the background

We proceed as in appendix B.2.1. Depending on $\bar{b}^{1:M}$, we apply one of the choices:

- If there is no active step, we proceed identically to appendix B.2.1.
- If there is at least one active step, instead of updating only $c$ we update $c$ jointly with the intensity $\bar{h}^{\text{last}}$ of the last active step. In this case, we maintain the same stochastic proposals for $c$ and adopt deterministic adjustments for $\bar{h}^{\text{last}}$. Our adjustments increase $\bar{h}^{\text{last}}$ by the same amount as $c$ decreases and, vice versa, decrease $\bar{h}^{\text{last}}$ by the same amount as $c$ increases. This is achieved by an adjustment function of the form:

$$f(c^{\text{prop}}, c^{\text{old}}) = c^{\text{old}} - c^{\text{prop}}.$$

This linear adjustment allows the stimuli before $\bar{h}^{\text{last}}$ to remain unchanged, in this way guiding the sampler towards high-probability regions of the parameter space.

#### B.3.2 Sampling the step intensities

We proceed as in appendix B.2.2; however, in the case $\bar{b}^m = 1$, we apply one of the choices:

- If there is no earlier active step, we proceed identically to appendix B.2.2.
- If there is at least one earlier active step, instead of updating only $\bar{h}^m$, we update $\bar{h}^m$ jointly with the intensity $\bar{h}^{\text{prev}}$ of the previous active step. In this case, we maintain the same stochastic proposals for $\bar{h}^m$ and adopt deterministic adjustments for $\bar{h}^{\text{prev}}$. Our adjustments increase $\bar{h}^{\text{prev}}$ by the same amount as $\bar{h}^m$ decreases and, vice versa, decrease $\bar{h}^{\text{prev}}$ by the same amount as $\bar{h}^m$ increases. This is achieved by an adjustment function of the form:

$$f\left((\bar{h}^m)^{\text{prop}}, (\bar{h}^m)^{\text{old}}\right) = (\bar{h}^m)^{\text{old}} - (\bar{h}^m)^{\text{prop}}.$$

This linear adjustment allows the stimuli before $\bar{h}^{\text{prev}}$ to remain unchanged, in this way guiding the sampler towards high-probability regions of the parameter space.

#### B.3.3 Sampling the step times

We proceed as in appendix B.2.3.

#### B.3.4 Sampling the step loads

We proceed as in appendix B.1.4 and appendix B.2.4; however, we choose, uniformly at random, to apply one of the following adjustments:

- If there is no earlier active step, we proceed identically to appendix B.1.4 and appendix B.2.4.

However, if there is at least one earlier active step, instead of updating only $\bar{b}^m$, we update $\bar{b}^m$ jointly with the intensity $\bar{h}^{\text{prev}}$ of the previous active step. In this case, we maintain the same stochastic proposals for $\bar{b}^m$ and adopt deterministic adjustments for $\bar{h}^{\text{prev}}$. Our adjustments decrease or increase $\bar{h}^{\text{prev}}$ by the same amount induced by activation or deactivation of $\bar{b}^m$. This is achieved by an adjustment function of the form:

$$f\left((\bar{b}^m)^{\text{prop}}, (\bar{b}^m)^{\text{old}}\right) = \begin{cases} -\bar{h}^m, & (\bar{b}^m)^{\text{prop}} = 1 \,\&\, (\bar{b}^m)^{\text{old}} = 0. \\ +\bar{h}^m, & (\bar{b}^m)^{\text{prop}} = 0 \,\&\, (\bar{b}^m)^{\text{old}} = 1. \end{cases}$$

This adjustment allows the stimuli before $\bar{b}^m$ to remain unchained, in this way guiding the sampler towards high-probability regions of the parameter space.

• If there is at least one later active step, instead of updating only $\bar{b}^m$, we update $\bar{b}^m$ jointly with the intensity $\bar{h}^{\text{next}}$ of the following active step. However, if there is no later active step, instead of updating only $\bar{b}^m$, we update $\bar{b}^m$ jointly with $c$. In both cases, we maintain the same stochastic proposals for $\bar{b}^m$ and adopt deterministic adjustments for $\bar{h}^{\text{next}}$ or $c$. Our adjustments increase or decrease $\bar{h}^{\text{next}}$ or $c$ by the same amount induced by activation or deactivation of $\bar{b}^m$. This is achieved by an adjustment function of the form:

$$f\left((\bar{b}^m)^{\text{prop}}, (\bar{b}^m)^{\text{old}}\right) = \begin{cases} -\bar{h}^m, & (\bar{b}^m)^{\text{prop}} = 1 \,\&\, (\bar{b}^m)^{\text{old}} = 0. \\ +\bar{h}^m, & (\bar{b}^m)^{\text{prop}} = 0 \,\&\, (\bar{b}^m)^{\text{old}} = 1. \end{cases}$$

This adjustment allows the stimuli after $\bar{b}^m$ to remain unchained, in this way guiding the sampler towards high-probability regions of the parameter space.

#### B.4 Sampler 4: A cutting-edge Gibbs scheme

Regardless of the deterministic adjustments, the sampler of appendix B.3 relies on the same stochastic proposals as the sampler of appendix B.2 and, although it can escape local maxima, it still contains adjustable parameters that need to be fine-tuned. Adaptation of the proposals during burn-in, in addition to requiring manual configuration, is based on heuristic rules [20, 21] that can lead to suboptimal mixing, as shown in RESULTS. Therefore, we propose *another novel sampler* that maintains multiplicative proposals such as the sampler in appendix B.2 and the ergodicity of the sampler in appendix B.3, but relaxes the sensitivity on fine-tuning. To achieve this, we reformulate our proposal mechanisms so that multiplicative sampling is performed via an elliptical slice sampler (ESS) instead of MH [14, 15, 17].

Before we describe the fine details of our sampling procedures, we present an overview of our novel sampling mechanism. For this, we start with a generic target $\pi(x)$ for a scalar variable $x$ attaining strictly positive values and complete it with an auxiliary variable

$$y \sim \text{Normal}(0, \rho^2).$$

With this completion, our aim is to introduce a normal variate that enables the application of ESS. Following the completion, our target becomes

$$p(x, y) = \pi(x)\text{Normal}\left(y; 0, \rho^2\right).$$

To sample the completed target, we apply four successive stages:

- Obtain $y$ by sampling its conditional $p(y|x)$. Due to our choices, this conditional reduces to Normal $(y; 0, \rho^2)$, which is sampled directly.
- Transform the completed target from $p(x, y)$ to

  $$P(X, Y) = e^{-Y}\pi\left(Xe^{-Y}\right)\text{Normal}\left(Y; 0, \rho^2\right),$$

  which is obtained under a bivariate transformation of random variables [18]. Specifically, this transformation is defined by

  $$\begin{pmatrix} x \\ y \end{pmatrix} \mapsto \begin{pmatrix} X \\ Y \end{pmatrix} = \begin{pmatrix} xe^{+y} \\ y \end{pmatrix}.$$

  With this transformation, our aim is to enable indirect updates of $x$ while resampling $Y$.

- Update $Y$ in the transformed target by sampling from its conditional $P(Y|X)$. Due to the form of the transformed target, this conditional is compatible with ESS which is our sampler of choice [14].
- Obtain the updated $x, y$ in the original completed target by the inverse transformation

$$\begin{pmatrix} X \\ Y \end{pmatrix} \mapsto \begin{pmatrix} x \\ y \end{pmatrix} = \begin{pmatrix} Xe^{-Y} \\ Y \end{pmatrix}.$$

  Because $y$ is sampled directly at the beginning of our scheme, at this stage the inverse transform only needs to recover $x$.

Our entire sampling mechanism, starting with some $x^{\text{old}}$ and leading to an updated $x^{\text{new}}$, is summarized in the sequence

$$x^{\text{old}} \xRightarrow[y\sim\text{Normal}(0,\rho^2)]{\text{direct sampling}} \begin{pmatrix} x^{\text{old}} \\ y^{\text{old}} \end{pmatrix} \xrightarrow[(x,y)\mapsto(X,Y)]{\text{transform}} \begin{pmatrix} X^{\text{old}} \\ Y^{\text{old}} \end{pmatrix} \xRightarrow[Y\sim P(Y|X)]{\text{ESS}} \begin{pmatrix} X^{\text{old}} \\ Y^{\text{new}} \end{pmatrix} \xrightarrow[(X,Y)\mapsto(x,y)]{\text{transform back}} \begin{pmatrix} x^{\text{new}} \\ y^{\text{new}} \end{pmatrix} \to x^{\text{new}}$$

where double arrows indicate random sampling and single arrows indicate deterministic transformations.

##### B.4.1 Sampling the background

We proceed as in appendix B.3.1; however, instead of performing resampling of $c$ with MH, we apply ESS as described above.

##### B.4.2 Sampling the step intensities

We proceed as in appendix B.3.2; however, instead of performing resampling of $\bar{h}^m$ with MH, we apply ESS as described above.

##### B.4.3 Sampling the step times

We proceed as in appendix B.2.3 and appendix B.3.3.

##### B.4.4 Sampling the step loads

We proceed as in appendix B.3.3.

### C Measurements and signal reduction

Measurements $w_n$ and signals $U(t)$ in our formulation model unitful quantities. For instance, $w_n$ is reported on camera-specific units, commonly ADU or counts, and $U(t)$ is reported on photon rates, which in turn depend on the choice of time units. For this reason, to facilitate visual inspection of our data, we convert unitful measurements and signals to unitless ones, $\tilde{w}_n$ and $\tilde{U}(t)$, as following

$$\tilde{w}_n = \frac{w_n - \mu}{g}, \qquad \tilde{U}(t) = \tau_{\text{exp}} U(t).$$

Our conversions result to "reduced" measurements $\tilde{w}_{1:N}$ and signal $\tilde{U}(\cdot)$ attaining the same units and comparable values as with the stimuli $u_{1:N}$.

### D Data quality metrics

Following the notation of appendix A.1, the *signal-to-noise ratio* of the $\beta^{\text{th}}$ step in a trace is given by

$$\text{SNR}^{\beta}_{\text{stp}} = \frac{h^{\beta}_{\text{stp}}}{\sqrt{\frac{v}{g^2\tau^2_{\text{exp}}} + \frac{f}{\tau_{\text{exp}}}\left(c_{\text{bck}} + \sum_{\beta'=\beta+1}^{B} h^{\beta'}_{\text{stp}}\right) + \frac{1}{2}\frac{f}{\tau_{\text{exp}}} h^{\beta}_{\text{stp}}}}. \tag{S7}$$

This is the root-mean-square variant of the *discriminability index* [7, 32] applied on raw fluorescent measurements with the statistics of eq. (S3) across the time $t_{\text{stp}}^{\beta}$ of the $\beta^{\text{th}}$ event. Essentially, our $\text{SNR}_{\text{stp}}^{\beta}$ quantifies the change in measurements over a photobleaching event against the pooled average of the corresponding variances. This index has been shown to correlate well with the performance of analysis methods operating on data contaminated with multiplicative noise and reduces to the familiar $\Delta$mean/std in special cases of additive noise [7].

Similarly, the *duration-to-exposure ratio* of the $\beta^{\text{th}}$ step in a trace is given by

$$\text{DER}_{\text{stp}}^{\beta} = \frac{1}{\tau_{\text{exp}}} \int_{t_{\text{stp}}^{\beta-1}}^{t_{\text{stp}}^{\beta}} dt \sum_{n=1}^{N} \Big[ H\left(t - t_n - \tau_{\text{exp}}/2\right) - H\left(t - t_n + \tau_{\text{exp}}/2\right) \Big]. \tag{S8}$$

In this definition, the integral on the right-hand side evaluates to the total exposure period (i.e. full time the camera is responsive to incident photons) excluding dead periods (i.e. full time the camera is unresponsive to incident photons) between consecutive events. This is achieved assuming that $t_{\text{stp}}^{0} = t_1 - \tau_{\text{exp}}/2$ coincides with the onset of the first exposure and $H(\cdot)$ coincides with the ideal step response function in agreement with eq. (S1). According to our definition, each $\text{DER}_{\text{stp}}^{\beta}$ is a unitless number that takes values between $0$ and $N$ and indicates the *amount of time equivalent to exposure periods* that the $\beta^{\text{th}}$ step's signal lasts within the trace.

In view of eqs. (S7) and (S8), the SNR and DER of the entire trace can be characterized by the minimum and maximum across its steps. For clarity, these are given by

$$\text{SNR}^{\min} = \min \text{SNR}_{\text{stp}}^{1:B}, \qquad \text{DER}^{\min} = \min \text{DER}_{\text{stp}}^{1:B},$$
$$\text{SNR}^{\max} = \max \text{SNR}_{\text{stp}}^{1:B}, \qquad \text{DER}^{\max} = \min \text{DER}_{\text{stp}}^{1:B}.$$

### E Parameters, hyperparameters, and other settings

Tables S1 to S3 list our specific values for the various parameters, model hyperparameters, and sampler settings used in the generation of the data shown in RESULTS.

| | Parameter | Value | Units |
|---|---|---|---|
| read-out offset | $\mu$ | $4.35 \times 10^3$ | ADU |
| read-out variance | $v$ | $1.94 \times 10^3$ | $\text{ADU}^2$ |
| overall gain | $g$ | $0.64$ | ADU/photon |
| excess noise factor | $f$ | $2$ | photons |
| exposure time | $\tau_{\text{exp}}$ | 0.1 | s |
| dead time | $\tau_{\text{dead}}$ | $1.74 \times 10^{-3}$ | s |
| start time of experiment session | $t_{\min}$ | $t_1 - \tau_{\text{exp}}/2$ | - |
| end time of experiment session | $t_{\max}$ | $t_N + \tau_{\text{exp}}/2$ | - |
| number of frames | $N$ | 500 | unitless |

*Table S1: Table of parameters related to data acquisition.*

| | Parameter | Value | Units |
|---|---|---|---|
| number of model steps | $M$ | 25 | unitless |
| hyperparameter for $\bar{b}^m$ prior | $\gamma$ | 1 | unitless |
| shape parameter for $c$ prior | $\phi_{\mathrm{bck}}$ | 2 | unitless |
| scale parameter for $c$ prior | $c_{\mathrm{bck}}^{\mathrm{ref}}$ | $\tilde{w}_N/\tau_{\mathrm{exp}}$ | - |
| shape parameter for $\bar{h}^m$ prior | $\phi_{\mathrm{stp}}$ | 3 | unitless |
| scale parameter for $\bar{h}^m$ prior | $h_{\mathrm{stp}}^{\mathrm{ref}}$ | $(\tilde{w}_1 - \tilde{w}_N)/\tau_{\mathrm{exp}}$ | - |
| hyperparameter for $\bar{t}^m$ prior | $t_{\mathrm{stp}}^{\mathrm{min}}$ | $t_{\mathrm{min}}$ | - |
| hyperparameter for $\bar{t}^m$ prior | $t_{\mathrm{stp}}^{\mathrm{max}}$ | $t_{\mathrm{max}}$ | - |

*Table S2: Table of hyperparameters related to data analysis.*

| | Sampler Implementation | Parameter | Value | Units |
|---|---|---|---|---|
| additive r.w. for $c$ | appendix B.1.1 | $\sigma_c$ | 500 | photons/s |
| additive r.w. for $\bar{h}^m$ | appendix B.1.2 | $\sigma_h$ | 100 | photons/s |
| additive r.w. for $\bar{t}^m$ | appendix B.1.3 | $\sigma_t$ | 5 | s |
| multiplicative r.w. for $c$ | appendices B.2.1 and B.3.1 | $\alpha_c$ | 20 | unitless |
| multiplicative r.w. for $\bar{h}^m$ | appendices B.2.2 and B.3.2 | $\alpha_h$ | 20 | unitless |
| multiplicative slice sampler | appendices B.4.1 and B.4.2 | $\rho$ | 1 | unitless |

*Table S3: Table of parameters related to MCMC proposals.*

| Parameter | Min | Max | Units |
|---|---|---|---|
| $B$ | 1 | 15 | unitless |
| $c_{\mathrm{bck}}$ | $3.9 \times 10^5$ | $1.6 \times 10^6$ | photons/s |
| $h_{\mathrm{stp}}$ | $4.0 \times 10^3$ | $2.4 \times 10^4$ | photons/s |
| $t_{\mathrm{stp}}^{\beta}$ | $t_{\mathrm{min}}$ | $t_{\mathrm{max}}$ | - |

*Table S4: Parameter ranges used in scope assessment. The ranges of $c_{\mathrm{bck}}$ and $h_{\mathrm{stp}}$ were anchored around typical experimental values, for instance such as those in fig. 2, and extended well beyond them by 0.2x and 2x to allow assessment of the performance of our methods under both typical and extreme fluorescence scenarios.*

### F Sensitivity analysis

Table S5 shows characteristic results that quantify the dependence of our methods on selecting model hyperparameters.

| | Sensitivity on $c_{\text{bck}}^{\text{ref}}$ | | | Sensitivity on $h_{\text{stp}}^{\text{ref}}$ | | | |
|---|---|---|---|---|---|---|---|
| Parameter | 0.5x | Baseline | 2x | 0.5x | Baseline | 2x | Units |
| $t^1$ | $-0.01$ | $-0.01$ | $0.00$ | $0.01$ | $-0.01$ | $-0.01$ | s |
| $t^2$ | $-0.01$ | $0.01$ | $-0.01$ | $-0.03$ | $0.01$ | $-0.01$ | s |
| $t^3$ | $-0.01$ | $-0.01$ | $-0.03$ | $0.01$ | $-0.01$ | $-0.01$ | s |
| $h^1$ | $-1.02$ | $1.25$ | $-0.26$ | $-1.07$ | $1.25$ | $-0.01$ | photons/ms |
| $h^2$ | $0.91$ | $-0.20$ | $-0.11$ | $0.03$ | $-0.20$ | $0.17$ | photons/ms |
| $h^3$ | $-0.83$ | $-0.10$ | $-0.16$ | $-0.25$ | $-0.10$ | $0.18$ | photons/ms |
| $c$ | $0.02$ | $-0.07$ | $0.00$ | $0.08$ | $-0.07$ | $0.03$ | photons/ms |

*Table S5: Local sensitivity analysis for hyperparameters $c_{\text{bck}}^{\text{ref}}$ and $h_{\text{stp}}^{\text{ref}}$. For each choice we calculate the bias (difference between posterior mean and ground truth) using a baseline value (as in table S2), that we also half (0.5x) and double (2x).*

### G Additional Figures

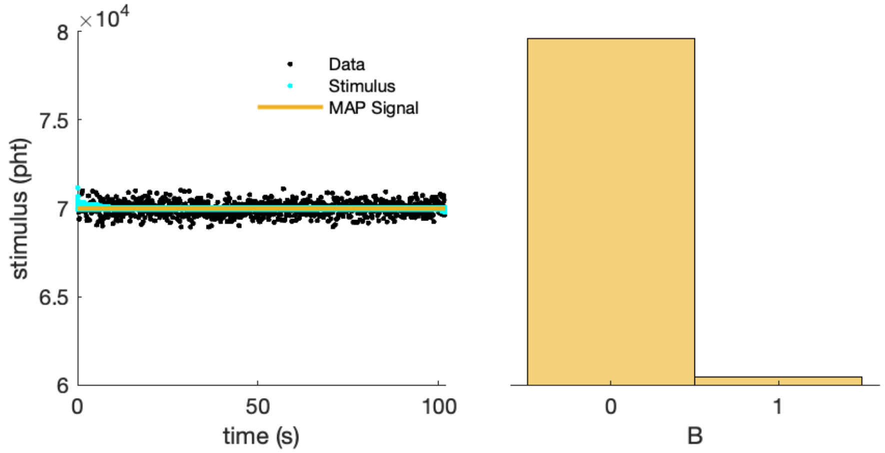

*Figure S2: Example of a trace from the dark channel containing no fluorophores. On the left panel we plot the data, stimulus, and MAP signal; while on the right we show a histogram of the sampled total number of active steps. As expected, our methods identify the presence of no steps and reconstruct the right signal with high confidence.*

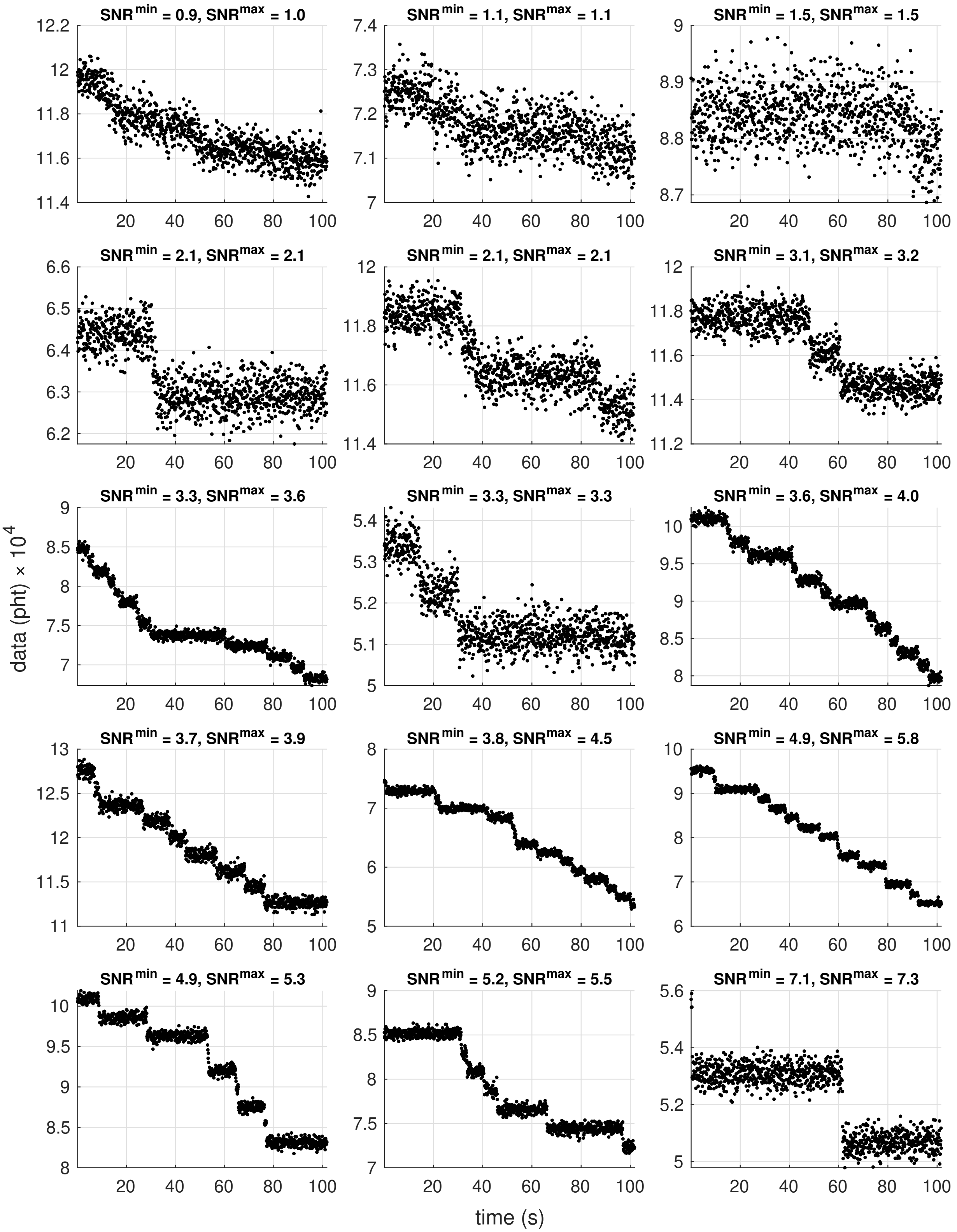

*Figure S3: Example synthetic data used in scope assessment along with their corresponding SNR*$^{\mathrm{min}}$ *and SNR*$^{\mathrm{max}}$*. To increase clarity, the vertical axes are* not *scaled consistently across panels; instead, they are adapted to their respective data.*

[6] Andrew Gelman, John B Carlin, Hal S Stern, and Donald B Rubin. *Bayesian data analysis*. Chapman and Hall/CRC, 2020.

[18] Steve Presse and Ioannis Sgouralis. *Data Modeling for the Sciences: Applications, Basics, Computations*. Cambridge University Press, 2023.

[19] Andreas Raue, Clemens Kreutz, Fabian Joachim Theis, and Jens Timmer. Joining forces of bayesian and frequentist methodology: a study for inference in the presence of non-identifiability. *Philosophical Transactions of the Royal Society A: Mathematical, Physical and Engineering Sciences*, 371(1984):20110544, 2013.

[20] Christian P Robert, George Casella, and George Casella. *Introducing monte carlo methods with r*, volume 18. Springer, 2010.
